# Complete zooplankton size spectra re-constructed from « in situ » imaging and Multinet data in the global ocean

**DOI:** 10.1101/2023.06.29.547051

**Authors:** Yawouvi Dodji Soviadan, Mathilde Dugenne, Laetitia Drago, Tristan Biard, Emilia Trudnowska, Fabien Lombard, Jean-Baptiste Romagnan, Jean-Louis Jamet, Rainer Kiko, Gabriel Gorsky, Lars Stemmann

## Abstract

Plankton size spectra are important indicators of the ecosystem state, as they illustrate the quantity of organisms available for higher marine food web and reflect multiple size-dependent processes. Yet, such measurements are typically biased by the available sampling methods, either disrupting fragile organisms or lacking good resolution (in size and/or time and space). In this study, we combined two of the most common approaches to measure zooplankton Normalized Biomass/Biovolume Size Spectra (NBSS) to calculate a complete zooplankton distribution for organisms larger than 1 mm. The reconstructed NBSS slopes appeared steeper and closer to those measured by the UVP5 (+7.6%) and flatter than those of the Multinet (- 20%) particularly in tropics and temperate latitudes. The overall gain in polar biomass was relatively small for reconstructed biomass compared to bulk estimates from Multinet (+0.24 mgC/m3 or +4.25%) and high from the UVP5 (+2.0 mgC/m3 or +53%). In contrast, in the tropical and temperate ecosystems, the gain in biomass was small for UVP5 (+0.67 mgC/m3 or +30.44% and +0.74 mgC/m3 or +19.59% respectively) and high for Multinet (+1.66 mgC/m3 or +136% and +3.4 mgC/m3 or +309% respectively). Given these differences, we suggest here to combine *in situ* imaging sensors and net data in any comprehensive study exploring key living players in the ocean ecosystem and their contributions to the biological pump.

## Introduction

Plankton are ubiquitous in the ocean and play important roles in trophic webs and biogeochemical cycles (Longhurst and Glen Harrison, 1989; Turner, 2002, 2015; Steinberg and Landry, 2017; Boyd *et al*., 2019). In particular, heterotrophic zooplankton are essential drivers of the transfer of primary production to higher trophic layers (Turner, 2004; Frederiksen *et al*., 2006), or to deep layers where carbon may be sequestered and stored for long periods of time (Cavan *et al*., 2017; Boyd *et al*., 2019). Their size range spans several orders of magnitude from single-cell eukaryotes (*i.e*. protists) to large jellyfish, with abundances decreasing exponentially with the size. This property is encapsulated in the Normalized Biovolume Size Spectrum (NBSS hereafter) approach, commonly used by scientists to study plankton and their size distributions. Through systematic measurements of organism’ abundances in increasing size classes, ecologists have shown that the shape of the NBSS varied temporally and spatially depending on the ecosystem functioning, and could thus be used as an indicator of the state of the ecosystem (Zhou, 2006; Frangoulis *et al*., 2010; Petchey and Belgrano, 2010; Gómez- Canchong *et al*., 2013). Indeed, the intercept of the NBSS can be used as a proxy of the biomass available at the base of the food web, while its slope indicates how biomass is transferred across sizes, through biological (e.g. predation, growth, remineralization, re-packaging) or physical processes (e.g. vertical or horizontal entrainments, sinking).

The overall estimates of zooplankton NBSS show that more information on plankton distribution on different spatio-temporal scales are required to accurately understand their ecology and contribution to pelagic processes under present and future environmental forcings (Dai *et al*., 2016; Ljungström *et al*., 2020). The traditional method to collect and study zooplankton at discrete locations has been plankton nets for almost two centuries, yet a significant fraction of the organisms may be under-sampled with this technique due to extrusion through the net mesh, entanglement within the net, and destruction of fragile forms such as gelatinous zooplankton (Bathmann *et al*., 2001; Gallienne and Robins. D. B, 2001; Warren *et al*., 2001; Remsen *et al*., 2004). Typically, these methods overlooked the importance of several zooplankton groups, such as rhizarians (Dennett *et al*., 2002; Remsen *et al*., 2004; Stemmann *et al*., 2008; Biard *et al*., 2016), or annelids (Christiansen *et al*., 2018). To address these limitations, non-destructive cameras have been developed to identify and quantify the abundance, size, and derived biomass of zooplankton *in situ* (Benfield *et al*., 2007; Picheral *et al*., 2010, 2022; Stemmann and Boss, 2012; Lombard *et al*., 2019). While improvements in Artificial Intelligence and Machine Learning are needed for these new camera devices to be broadly adopted by zooplankton researchers, particularly taxonomists (Irisson *et al*., 2022), they provide the increased spatial and temporal resolution needed to study the coupling between physical processes and zooplankton distributions and for modeling zooplankton community and trophodynamics (Lombard *et al*., 2019; Drago *et al*., 2022; Giering *et al*., 2022; Soviadan *et al*., 2022).

In the recent survey by (Giering *et al*., 2022), a panel of scientists has recommended to retain net sampling and traditional taxonomy in the future, in combination with a period of overlapping use of both sampling systems to ensure the continuity and replacement of physical sampling by *in situ* imaging in zooplankton monitoring programs. Despite such recommendations, the comparison of results from nets and imaging cameras at global scale of the open ocean (tropical, temperate and polar systems) in the 0-1000m water column is still rare. The lack of systematic and consistent analysis impairs a global assessment of zooplankton composition and biomass. The present study uses a combination of observations from Multinet and Underwater Vision Profiler 5 (UVP5) in 57 stations located in all oceans and 5 depth layers in the upper kilometer, to reconstruct a complete representation of zooplankton NBSS, and derived biomass, in the large size fractions (> 1 mm). We then compare new NBSS estimates to those by each method to evaluate the differences and similarities. We discuss the strength and limitations of each approach to describe complete zooplankton communities.

## Materials and methods

### Zooplankton sampling and imaging

The Tara Oceans expedition (Karsenti *et al*., 2011) and Tara Polar Circle took place between 2009 and 2013. Among the 210 stations crossed during the expedition in the different ocean basins, 57 stations (Fig. 1.) were sampled with both a Multinet (Roullier *et al*., 2014; Pesant *et al*., 2015) and an Underwater Vision Profiler 5 (Picheral *et al*., 2010). Sampling covered a variety of ecosystems ranging from oligotrophic to eutrophic. The UVP was mounted on the vertically profiling system and recorded and quantified the size and abundance of specific groups of zooplankton >600 m. Concordant measurements of zooplankton size and abundance were thus obtained from these two different approaches. The sampling and processing steps are described in detail below.

**Figure 1:**
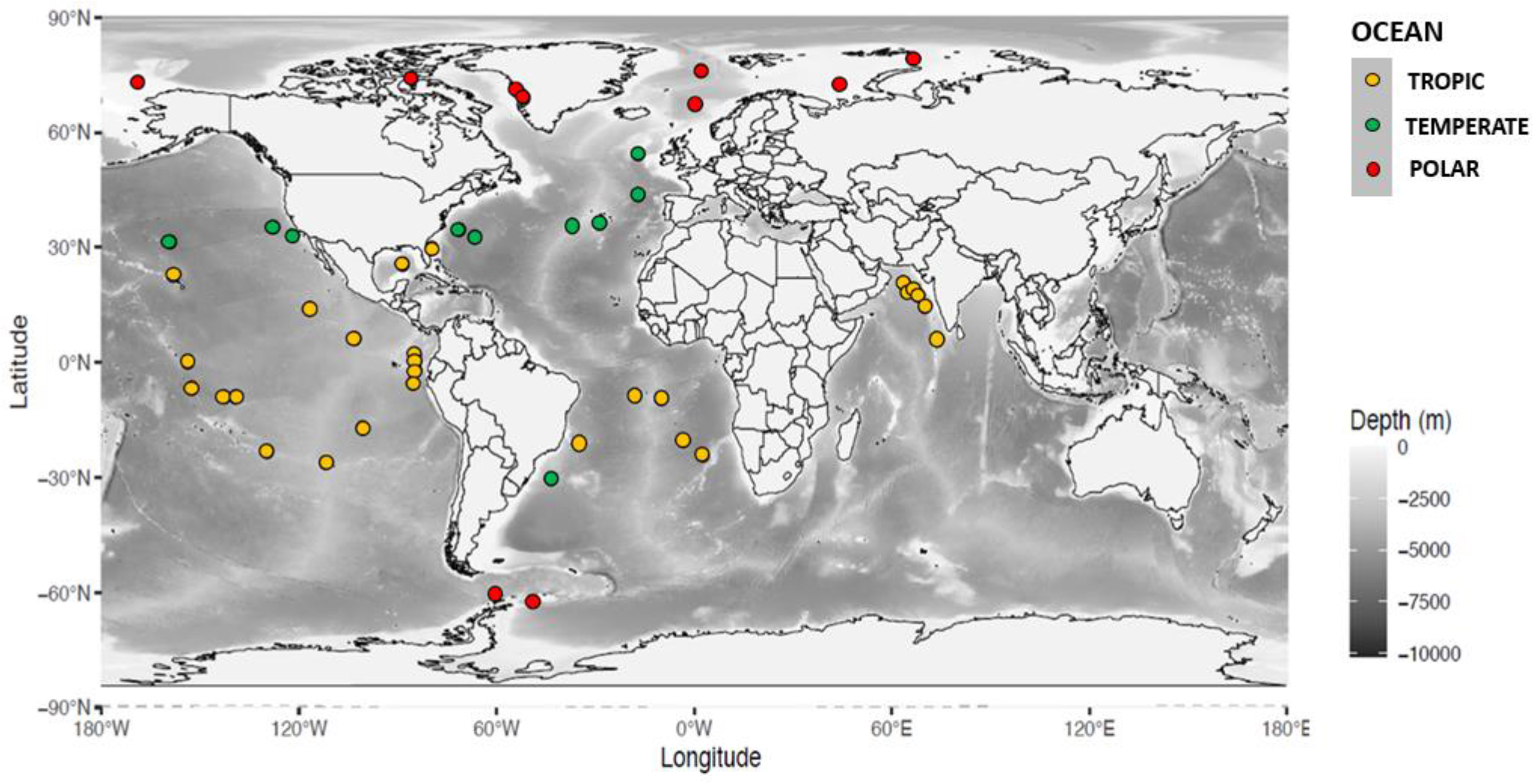
Location of the 57 stations sampled with UVP5 and Multinet systems, grouped by latitudinal regions coloured in yellow (tropic: 0-30°), green (temperate: 30-60°) and red (polar: > 60°), superimposed on global bathymetry.

The Multinet, deployed vertically, was composed of five sequential plankton nets equipped with a 300 µm mesh and an aperture of 0.25 m^2^. It was equipped with a flow meter to estimate the sampled volume which ranged from 5 to 502 m^3^ (median value of 113 m^3^). All five nets sampled plankton at five different depth layers distributed between the surface and 1000 m based on an adaptive strategy depending on observed physical or biological features such as the deep chlorophyll maximum (see rationale in (Soviadan *et al*., 2022)) across the 57 global stations. Collected samples were preserved in a solution of buffered formaldehyde (4% final concentration). In the laboratory, the samples were rinsed and fractionated with a Motoda box. The final fraction was scanned with the ZooScan system and processed with Zooprocess software which allowed a rapid and time-efficient analysis of the plankton samples in a digital format that can be easily stored before further processing (Gorsky *et al*., 2010).

Also, *in situ* UVP5 profiles provided automatic information on all particles larger than 100µm detected by the sensor, in addition to specific information on large zooplankton groups (area >30 pixels in size, approximately equivalent spherical diameter > 600µm) that could be taxonomically identified. The UVP5 recorded a maximum of one image of about 1 liter volume every 5 cm for a 1m s^−1^ lowering speed, resulting in significantly lower sampled volume compared to the Multinet, ranging from 0.033 to 21.23 m^3^ (median value of 4.32 m^3^). Therefore, all profiles of the UVP5 obtained at a given station were merged and the counts of all automatically detected zooplankton were integrated over the five depth layers corresponding to the discrete depth layers sampled by the nets at the same station. With this data aggregation strategy, we cumulated a sampled volume greater than 40 m^3^ for each calculated NBSS which ensures a good representation of rare organisms in the water column.

The datasets were grouped in three latitudinal bands (inter-tropical, temperate and polar), and three depth layers (0-200, 200-500, 500-1000m) to explore NBSS shapes and plankton community compositions across the globe. Here, we consider the tropical and temperate bands to be larger than their theoretical geographic limits, spanning from 30°N to 30°S latitude and up to 60° S and N.

In total, the Multinet samples comprised nearly 400,000 images of organisms whereas the UVP5 image collection consisted of 769,297 images (including living and non-living particles), of which 5% (n=37,000) were of zooplankton. Both sets of images were uploaded to Ecotaxa (http://ecotaxa.obs-vlfr.fr), an online collaborative platform allowing visual validation of the taxonomic classification of each organism predicted with a semi-automatic classifier integrated to the Ecotaxa web application. The images were validated into different groups for each instrument. For the ZooScan, the high definition images allowed to identify down to the family (and sometimes down to genus) rank, as presented in (Soviadan *et al*., 2022). However, due to the lower definition of the UVP5 images and the smaller cumulative volume, we restricted our NBSS estimates to 8 categories described in Table I.

**Table I.**
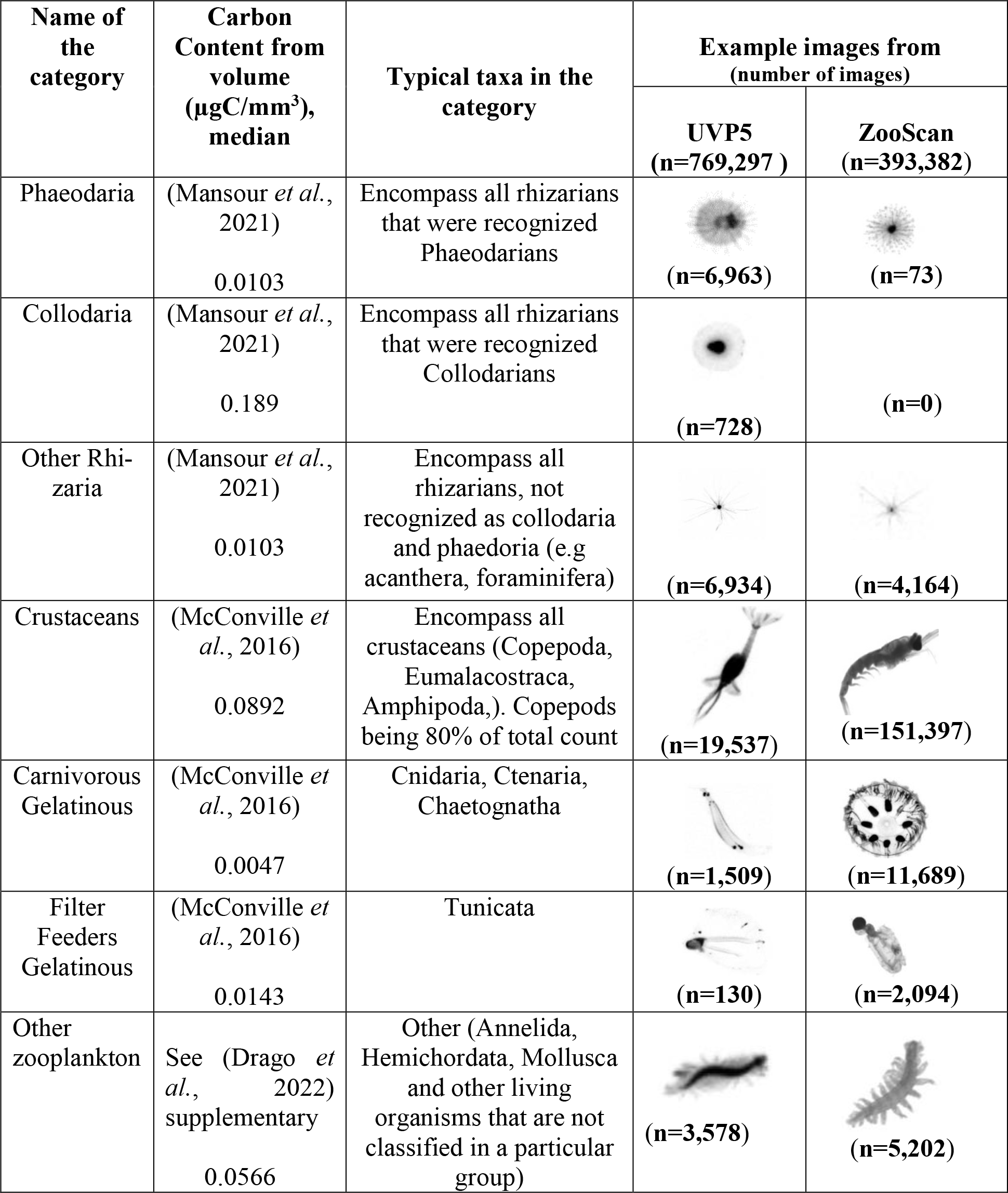
: List and examples of images of zooplankton taxa identified in the present study and conversion factors from biovolume to carbon content.

| Name of the category | Carbon Content from volume ($\mu\text{gC}/\text{mm}^3$), median | Typical taxa in the category | Example images from (number of images) | |
| --- | --- | --- | --- | --- |
|  |  |  | UVP5 (n=769,297 ) | ZooScan (n=393,382) |
| Phaeodaria | (Mansour *et al.*, 2021)<br>0.0103 | Encompass all rhizarians that were recognized Phaeodarians | (n=6,963) | (n=73) |
| Collodaria | (Mansour *et al.*, 2021)<br>0.189 | Encompass all rhizarians that were recognized Collodarians | (n=728) | (n=0) |
| Other Rhizaria | (Mansour *et al.*, 2021)<br>0.0103 | Encompass all rhizarians, not recognized as collodaria and phaeodaria (e.g acanthera, foraminifera) | (n=6,934) | (n=4,164) |
| Crustaceans | (McConville *et al.*, 2016)<br>0.0892 | Encompass all crustaceans (Copepoda, Eumalacostraca, Amphipoda,). Copepods being 80% of total count | (n=19,537) | (n=151,397) |
| Carnivorous Gelatinous | (McConville *et al.*, 2016)<br>0.0047 | Cnidaria, Ctenaria, Chaetognatha | (n=1,509) | (n=11,689) |
| Filter Feeders Gelatinous | (McConville *et al.*, 2016)<br>0.0143 | Tunicata | (n=130) | (n=2,094) |
| Other zooplankton | See (Drago *et al.*, 2022) supplementary<br>0.0566 | Other (Annelida, Hemichordata, Mollusca and other living organisms that are not classified in a particular group) | (n=3,578) | (n=5,202) |

Once the zooplankton images were validated on Ecotaxa, the concentration and morphometric measurements of each living organism were extracted for every station and net. The biovolume was estimated using the minor and major ellipsoidal axis provided by Zooprocess assuming an ellipsoidal shape (Gorsky *et al*., 2010). The derived equivalent spherical diameter of individual zooplankton varied from 0.25 to 20 mm for the Multinet images and from 0.6 mm to 20 mm for the UVP5 images. To extend our comparisons to other studies, plankton biovolume was converted into dry mass then C biomass using taxa-specific relationship between size and mass found in the literature (Table supplementary I).

### Net- and UVP-based plankton size distribution and NBSS estimates

Plankton size distributions were estimated using the broadly-used NBSS which was initially developed for zooplankton (Denman and Platt, 1978). They were obtained by sorting the individual biovolume of each organism in increasing logarithmically-spaced size classes defined by [log(X_n_); log(X_n+1_)], with equal distance: log(X_n+1_) − log(X_n_) = log(k), the constant k is 2^1/4^ . In general, all NBSS present a mode in the size spectrum at the lower size range, reflecting the minimum size of efficient detection and processing by imaging system, while high variability in the large size range reflects a relatively small sampled volume for that size range (Stemmann and Boss, 2012). In our study, we considered a maximum of 20 size classes to calculate the NBSS. Smaller organisms were likely underestimated because of the mesh size or the threshold used to process the raw images, hence we determined the smallest size class using the approach described by (García-Comas *et al*., 2014), which corresponds to the size class where the first maximum NBSS was observed in the net samples (see Table Supplementary I).

NBSS was calculated by dividing the summed biovolume [Σbiovolume (mm^3^)] in each size class by the cumulative sampled volume (m^3^), and further dividing this ratio by the width of each size class interval [Δbiovolume (mm^3^)]. Using these size spectra, we extracted the NBSS slope which is obtained by simple log-linear regression on the linear part of the size spectrum: 1.46-7.34 mm for the Multinet NBSS and 1.47-7.45 mm for the UVP5 NBSS. Since these linear portions differed between the two approaches, we chose the lower threshold of 1 mm to effectively compare biomass in this study. This value corresponds to the size where both median NBSS values overlapped, and were similar to the value of 0.934 mm used by (Barth and Stone, 2022) to compare these two methods during 5-day cruises of the Bermuda Atlantic Time-series Study. The upper threshold has been set at 8 mm. (Drago *et al*., 2022) also estimated the total biomass of zooplankton within a size range of 1-50 mm from a global UVP5 dataset. Hereafter, we refer to NBSS estimates derived from zooplankton collected by the Multinet or imaged by the UVP5 as NBSS_Zmtn and NBSS_Zuvp, respectively.

### Reconstruction of zooplankton NBSS and comparison with the Mulinet and the UVP estimates

To construct a realistic representation of NBSS estimates, we looked at individual taxonomic group results and identified the paired observations where both UVP5- and Multinet- based NBSS had finite values, with the exception of groups of rhizarians (*e.g*. Collodaria, Phaeodaria and Other Rhizaria) which had no finite values in Multinet NBSS in the >1mm size range (Fig. 2.). For these groups, we only used the UVP5-based NBSS. We note that another group of rhizarians was above the detection limit in several Multinet samples but their size estimates were always below 1 mm in Multinet samples. This group appeared strongly correlated to the UVP5 larger (> 1 mm) “other rhizarians” (see Figure 2 Supplementary). Given this correlation and the taxonomic affiliation of these images (classified as Foraminifera in the “other rhizarians”), we assumed that the same taxonomic group was imaged without their house in the Multinet samples and with their house by the UVP5, and thus also used the NBSS_Zuvp exclusively. We then selected the maximum of these non-null paired values (UVP5 and net) within each size class for each taxonomic group, and summed the values of the different taxonomic groups to obtain the bulk reconstructed NBSS estimates for all living.

**Figure 2:**
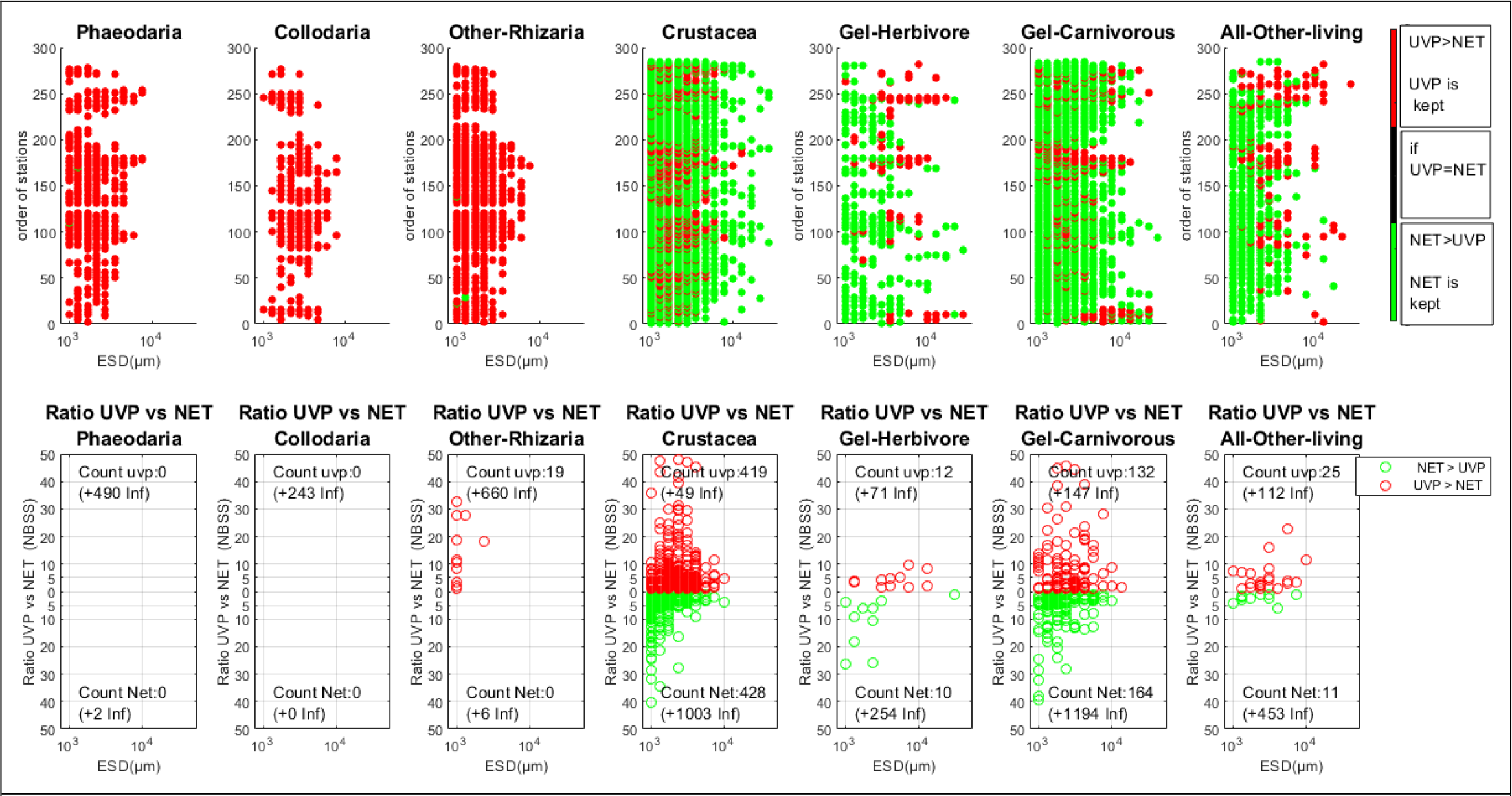
Illustration of the NBSS selection criteria to build the reconstructed NBSS. The top panel indicates NBSS estimates selection between UVP5 (red) or nets (green) based on the maximum value observed for individual taxonomic groups. The bottom panel shows the ratio of UVP/NET NBSS when the size class for paired of finite values. Note that Phaeodaria and Collodaria did not present finite NBSS estimates from Multinet samples.

Comparison of the reconstructed NBSS and NBSS_Zuvp or NBSS_Zmtn was done by computing the integrated biomass and slopes from all estimates. We used a non-parametric Kruskal-Wallis test to test for significant differences between slopes and integrated biomass, as well as the Spearman coefficient to test for correlations among variables.

## Results

### Reconstructed NBSS

To reconstruct the zooplankton NBSS in the 57 stations, we selected the maximum value observed in all paired finite NBSS estimates, in each size bin larger than 1 mm, and for each individual taxonomic group. The resulting selection is shown in Fig. 2. . Of all taxonomic groups, the rhizarians, Collodaria and Phaeodaria, were detected almost only in the UVP5 samples. For these groups, we picked UVP5-derived NBSS estimates. A third group, “Other- Rhizaria”, presented NBSS estimates above the detection limit for the Multinet and UVP5 samples, but with a very distinct size distribution for the two sampling methods (<1 mm in Multinet samples and >1 mm in UVP5 samples, see Supplementary Figure. 2b). The total concentration of this group across all paired samples observations were significantly correlated (r^2^=0.75, p-value=8.1x10^-56^, y=1.3x+2.24), suggesting that they might have been the same organisms (Figure 2a supplementary). Crustaceans, mainly represented by copepods, appeared well detected by Multinet in small (ESD<1mm) and large (ESD> 1mm) size classes, covering a larger size range than the UVP5. Intermediate size classes were generally better sampled with UVP5, with a median ratio (UVP/NET) of 2.77 and quartiles of 1.57 to 7.56 (n=419) when UVP>NET against a median ratio (NET/UVP) of 2.56 and quartiles of 1.57 to 4.57 (n=428) when NET>UVP. Similarly, gelatinous carnivorous organisms, were well detected with the UVP5 at intermediate size classes with a median ratio (UVP/NET) of 4.71 and quartiles of 1.89 to 12.37 (n=132) against a median ration (NET/UVP) of 3.33 and quartiles of 1.93 to 7.20 (n=164). Gelatinous herbivore and ‘Other’ living were mostly found in intermediate size classes by net and in large size classes by UVP5.

These differences in the number of organisms detected by the two approaches result in large offset of total C biomass. The total POC content (see Table I for conversion factors), derived from images biovolume estimates and the size-to-biomass factor (Figures 3 & 5 supplementary), showed that the contribution of rhizarians, gelatinous carnivores and filter feeders is not negligeable compared to other solid forms like crustaceans, despite their lower C content (Figure 4 supplementary). In tropical and temperate latitudinal bands, the former groups contributed more than 50% to the total plankton biomass in the epipelagic zone and about 40% in the mesopelagic zone. The rhizarians inhabiting the epipelagic layers were dominated by collodarians in the tropics and by the phaeodarians in temperate regions. Crustacean’s biomass was always higher than other zooplankton biomass in polar waters where both UVP5 and Multinet presented similar patterns. In this region, some phaeodarians appeared in deeper UVP5 casts, while they were absent from Multinet samples. Gelatinous carnivorous biomass appeared well observed with both approaches and always accompanied those of crustaceans.

**Figure 3:**
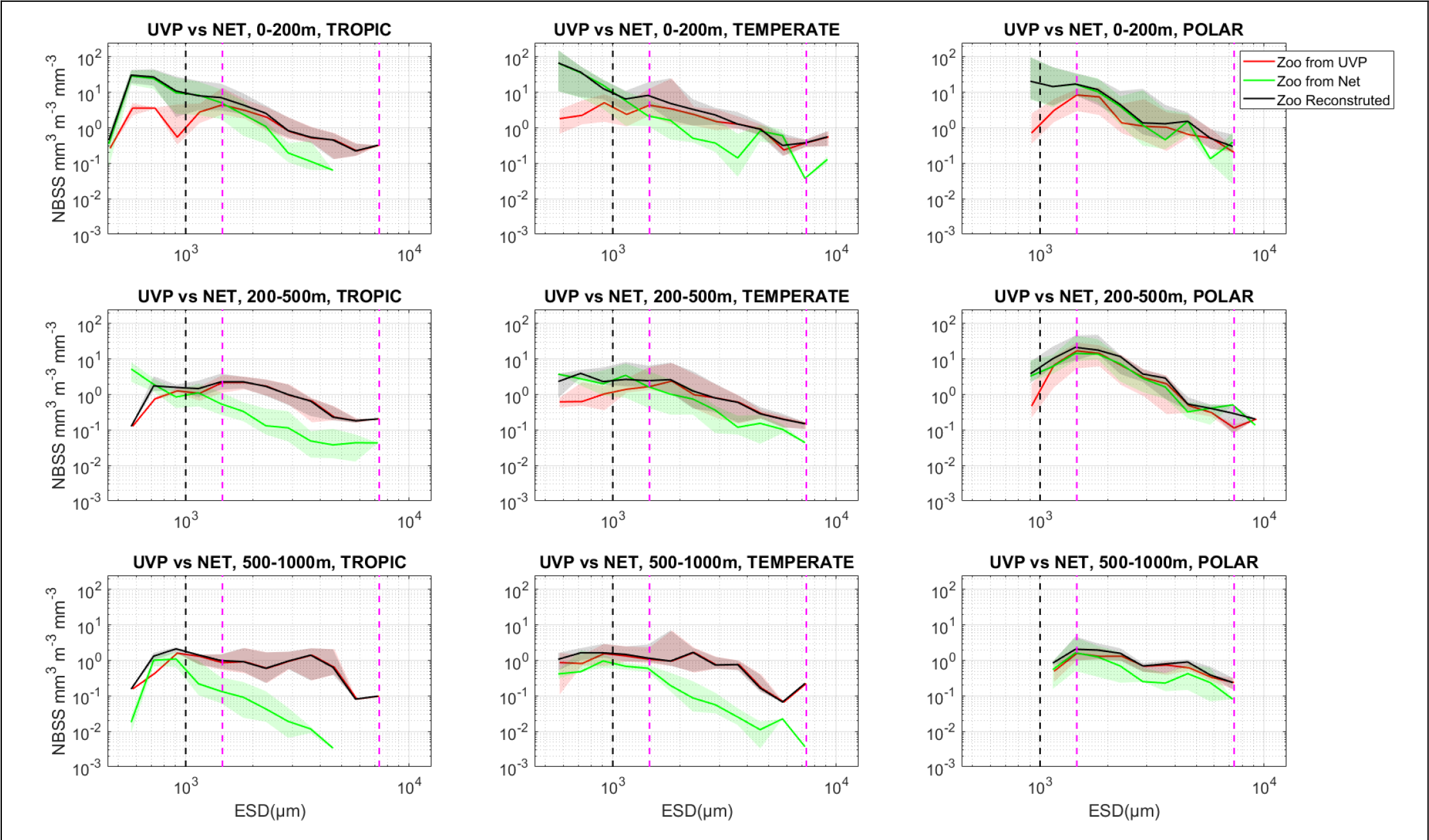
NBSS of all zooplankton detected by the UVP5 (red), nets (green), and the reconstructed NBSS (black, maximum values) in three depth layers (0-200, 200-500 and 500-1000m), and three latitudinal bands (0-30°, 30-60°, >60°). The shaded area corresponds to the 25 and 75 quartiles around the median of all NBSS observed at the 57 stations shown in Figure 1. The black dashed line corresponds to the 1mm while the marron dashed lines delimit the size range used to extract and compare the NBSS slopes.

**Figure 4:**
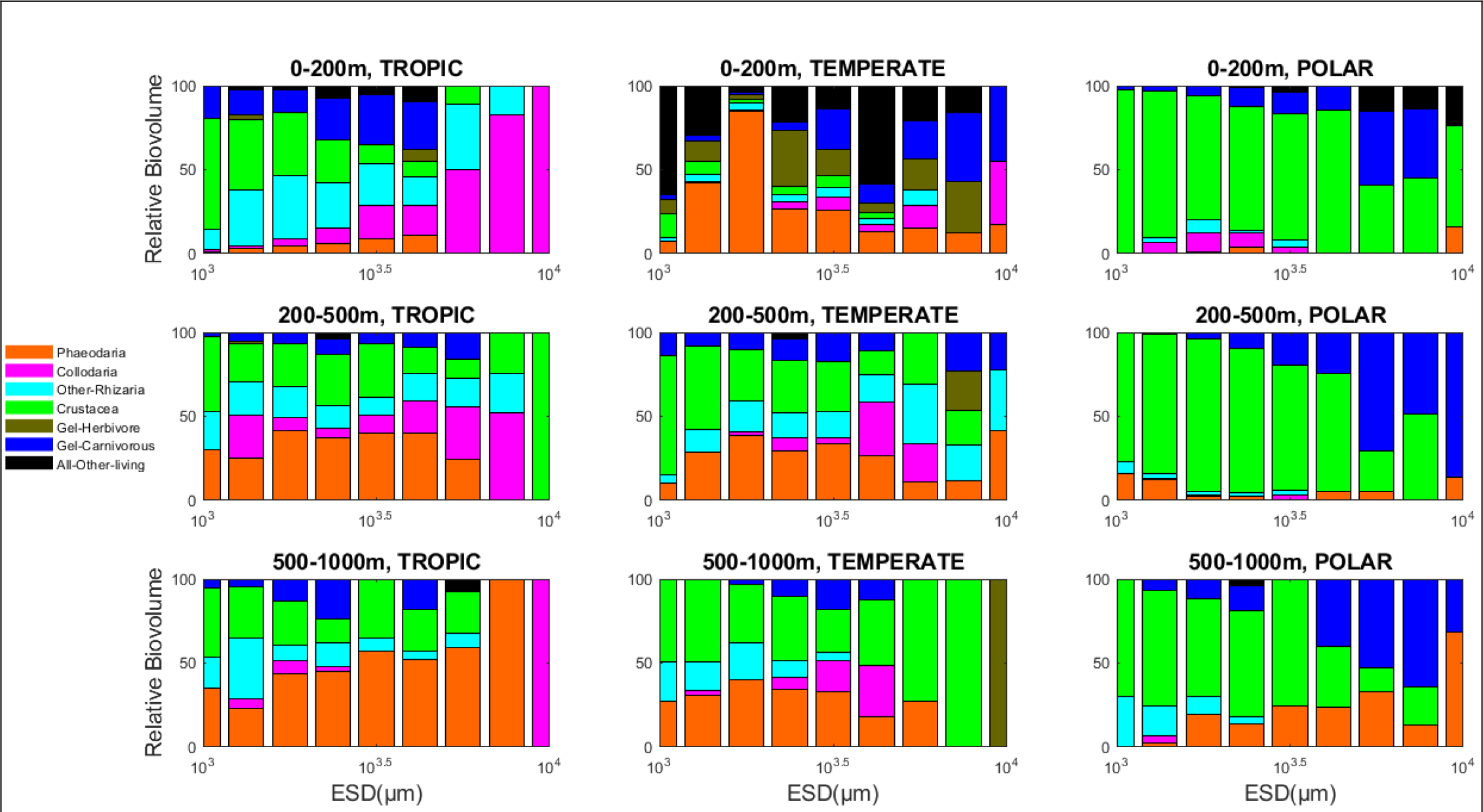
Relative contributions of different zooplankton taxa to the reconstructed NBSS (for plankton size from 1 to 8mm) in three depth layers (0-200, 200-500 and 500-1000 m) and three latitudinal bands (0°-30°, 30°-60°, >60).

### Zooplankton NBSS in three depth layers and latitudinal bands

Zooplankton NBSS estimated by UVP5 (NBSS_Zuvp) and Multinet (NBSS_Zmtn) show substantial differences notably at size larger than 1.21mm in ESD mainly in the tropical latitudinal band (Figure 7 supplementary). In general, NBSS_Zuvp estimates presented higher values than those derived from the Multinet at multiple depths and latitudes from tropical to temperate zones, for organisms larger than 1 mm. Multinet NBSS also declined more sharply with size, indicative of a steeper NBSS slopes, compared to the UVP5 estimates. The median ratio of NBSS_Zuvp to NBSS_Zmtn values, across all size classes and all taxonomic units, was 3.19 and quartiles of 1.70 to 9.26 at these locations. In contrast, the two NBSS estimates strongly overlapped at the surface and in the upper mesopelagic zone of the polar regions. For all latitudinal bands, integrated NBSS values, used as a proxy for total zooplankton abundances, showed a sharp decline in concentration with depth, with values in the 0-200 m depth (epipelagic) layer always greater than those in the mesopelagic layer (200-1000 m).

The reconstruction of zooplankton NBSS overlaps with UVP5- and Multinet-derived estimates (Fig. 3.). As expected, the reconstructed NBSS closely aligned to the upper envelop of the measured NBSS, and overall followed the UVP5-derived estimates more closely than the Multinet. The maximum concentration of all zooplankton taxonomic groups also declined with depth, with the epipelagic layer being on average 47 times higher than the upper mesopelagic layer, and 12 times higher than the lower mesopelagic layer.

### Relative contribution of different taxa to zooplankton reconstructed NBSS

The relative contribution of the different plankton categories (Fig. 4) identified from the reconstructed NBSS showed that the main contributors to the total biovolume were crustaceans and rhizarians (in most size fractions) and gelatinous carnivores in the largest size class. However, there was a large variability depending on the latitudinal bands and depth layers. The contribution of rhizarians to the total zooplankton biovolume in all size classes decreased from low latitudes to high latitudes in the epipelagic and mesopelagic layers. Inter-tropical samples showed that in the epipelagic layer, collodaria dominated in the larger size fraction while in the mesopelagic layer, phaeodarians dominated in the small size fraction. In temperate samples, phaeodarians and gelatinous (carnivorous and filter feeders) dominated the biovolume at the surface while copepods and phaeodarians were dominant in the mesopelagic. In the polar regions, crustaceans clearly dominated biomass and biovolume in almost all size classes excepted in large size classes of the lower mesopelagic layer, where carnivorous gelatinous and Phaeodarians were more abundant.

### Comparison of zooplankton NBSS slopes and total biomass by different methods

The NBSS slopes, and their variability around the median, varied by depth (Fig. 5.), with flatter, less variable, slopes in the mesopelagic zone and steeper slopes in the epipelagic layer. The slopes of the reconstructed NBSS in the tropical epipelagic layer presented a median value of -1, similar to the values reported in previous studies. The reconstructed NBSS slope appeared closer, albeit steeper, to that measured by the UVP5, and was systematically flatter than that of Multinet (p-value>0.05). No significant differences were found between slopes from UVP5 and the recontructed NBSS in the tropical and temperate epipelagic layer (p-value>0.05). Differences were not significant in the polar epipelagic layer for the three methods (p-values>0.05).

**Figure 5:**
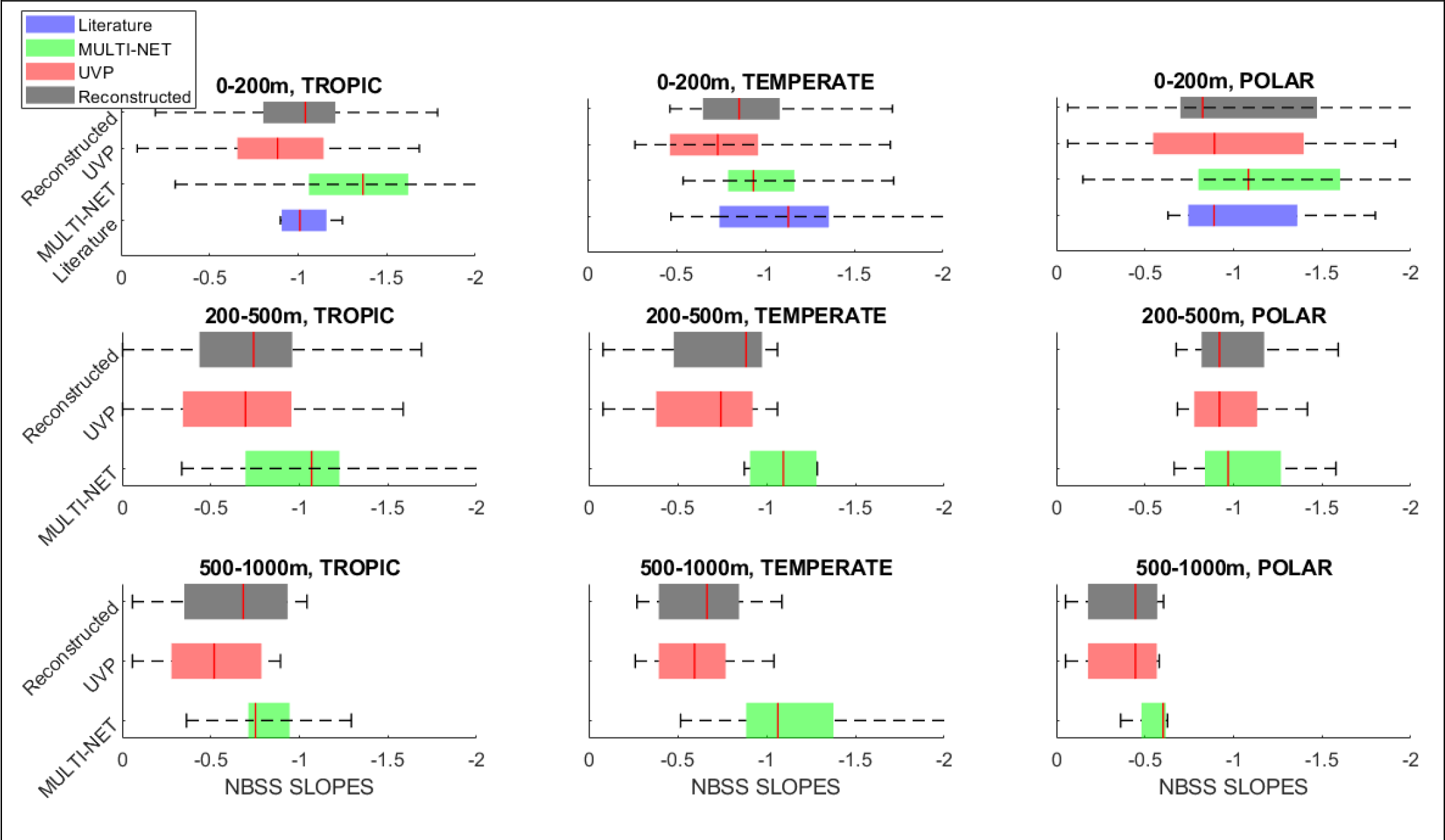
Comparison of mesozooplankton Normalized Biovolume Size Spectra (NBSS) slopes at different locations in three depth layers (0-200, 200-500 and 500-1000m), and three latitudinal bands (0-30°, 30-60°, >60°) for plankton size from 1 to 8mm. Our study data are from Multinet and UVP5. Statistical test is Kruskal-Wallis.

Our results showed that the reconstructed NBSS slopes were flatter than the Multinet (- 20%, correlation coefficient: 0.36 +/- 0.14, pvalue= 3.75 x10^−07^) and steeper than the ones of UVP5 (+7.6%, correlation coefficient: 0.93 +/- 0.02, pvalue=5.56 x10^−74^).

Like spectral slopes, the total reconstructed zooplankton biomass of organisms larger than 1 mm presented a general decrease with depth apart from the polar region where the biomass maximum was found in the upper mesopelagic (Fig. 6.). Total zooplankton biomass determined from the reconstructed NBSS showed a decrease from poles to the tropics. The reconstructed biomass was sytematically higher than that of the Multinet, with the largest differences observed in the tropical and temperate surface layers. In general the biomass of the reconstructed NBSS is closer to that of UVP5 in tropics and temperate oceans and almost similar to that of multinet in polar ocean.

**Figure 6:**
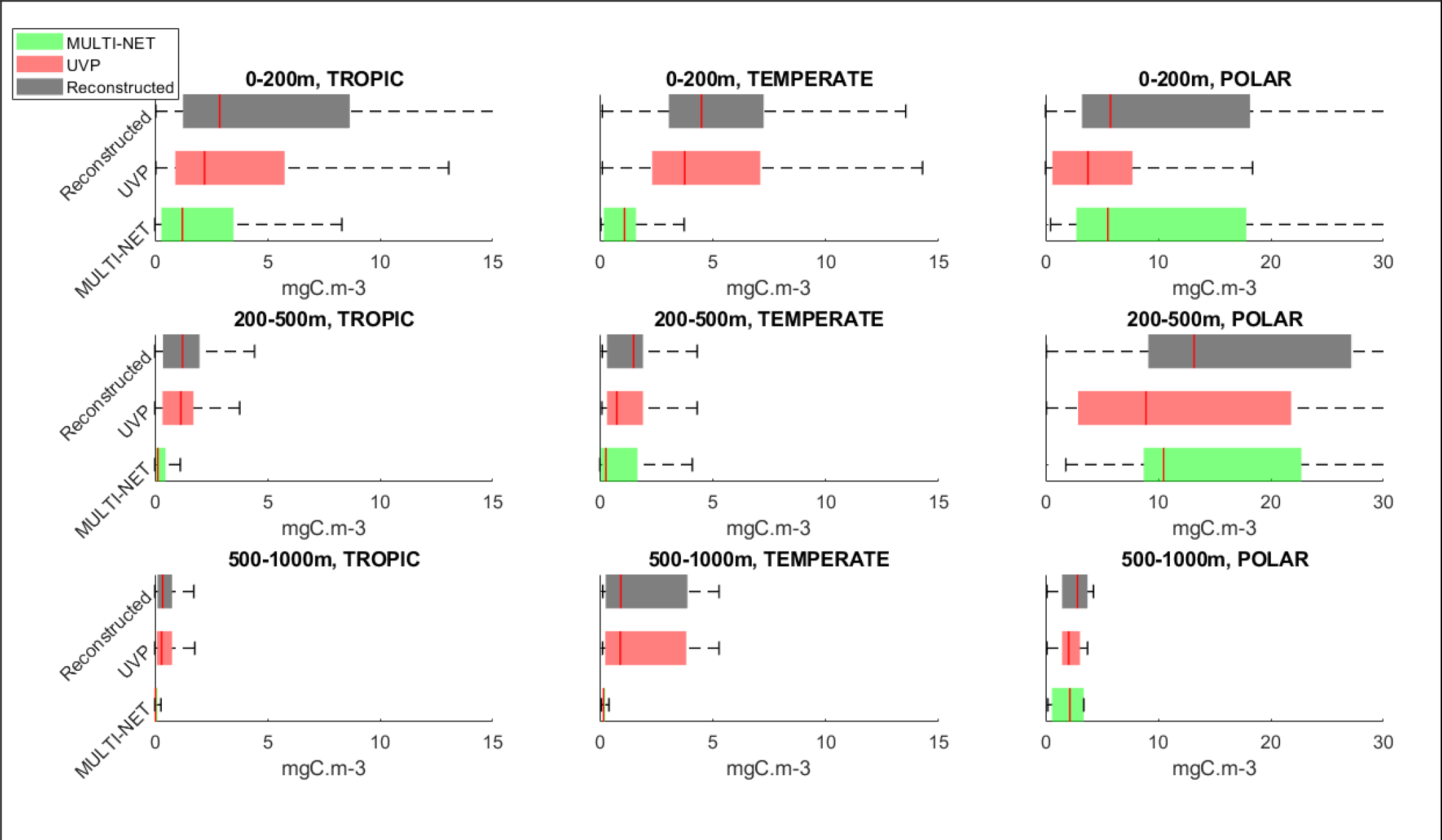
Comparison of mesozooplankton Biomass at different locations in three depth layers (0-200, 200-500 and 500-1000m), and three latitudinal bands (0-30°, 30-60°, >60°) for plankton size from 1 to 8mm. Our study data are from Multinet and UVP5. Statistical test is Kruskal-Wallis.

In the tropics, the absolute gain in biomass (Table Supplementary II) when comparing the reconstructed biomass and the Multinet decreased with depth, with +1.66 mgC m^-3^ (136%) in the surface, +0.81 mgC m^-3^ (615%) in the upper mesopelagic, and +0.27 mgC m^-3^ (900%) in the lower mesopelagic. In temperate ecosystems, the gains in biomass were significantly higher, with +3.4 mgC m^-^ (309%) in the surface, +1.23 mgC m^-3^ (440.1%) in the upper mesopelagic, and + 0.77 mgC m^-3^ (440.5%) in the lower mesopelagic. In polar ecosystems, gains were much lower, with +0.24 mgC m^-3^ (4.25%) in the surface, +2.71 mgC m^-3^ (26%) in the upper mesopelagic, and +0.68 mgC m^-3^ (31.66%) in the lower mesopelagic.

The same trends were observed when comparing the reconstructed biomass to that of the UVP5, although gains were generally more marginals. In the tropics, the average gain in biomass ranged from +0.04 mgC m^-3^ (14.2%) and +0.08 mgC m^-3^ (6.56%) in the lower and upper mesopelagic, to +0.67 mgC m^-3^ (30.44%) in the surface. In temperate ecosystems, reconstructed biomass was on average 19.59% (+0.74 mgC m^-3^), 2.71% (+0.024 mgC m^-3^), and 97% (+0.74 mgC m^-3^) higher in the epipelagic, lower and upper mesopelagic. In the same depth layers, polar ecosystems biomass gained 53% (+ 2.0 mgC m^-3^), 48% (+4.27 mgC m^-3^), and 38% (+0.78 mgC m^-3^).

## Discussion

Here, we provide a comprehensive dataset of zooplankton size distribution, taxonomic composition and total biomass based on individual measurements of body size and document the shape of the size spectrum on a global scale. Previous zooplankton studies have mostly reported NBSS estimates based on nets, or total biomass from UVP5 data (Forest *et al*., 2012; Biard *et al*., 2016; Drago *et al*., 2022). However, here we report new, complete, zooplankton size distribution estimates across the global ocean, using the combination of both net and UVP approaches. In the following sections, we compare our UVP5 and net datasets to other studies so far restricted to specific oceanic regions, we discuss discrepancies between each approach and finally highlight the novel understanding of the complete NBSS.

### Comparison of our global zooplankton dataset to existing case studies

We find that zooplankton communities collected with nets exhibit higher NBSS values towards the poles, a pattern that is largely driven by crustaceans. This pattern observed towards the poles, also described in (Brandão *et al*., 2021), is induced by the higher contributions of the large-size grazing Calanidae relative to the smaller omnivorous-carnivorous Cyclopoida and Poecilostomatoida (i.e. Oithonidae, Oncaeidae and Corycaeidae) that are more dominant in tropical regions (Brandão *et al*., 2021; Soviadan *et al*., 2022). It is also well known that zooplankton size decreases with increasing temperature and an increase in the relative contribution of small phytoplankton, while it increases with concentrations of oxygen, macronutrients, total phytoplankton biomass and the relative contribution of large phytoplankton (Brun *et al*., 2016; Brandão *et al*., 2021; Soviadan *et al*., 2022). This latitudinal trend is consistent with other observations of high Copepoda abundance or biomass in the Polar oceans (Hirche and Mumm, 1992; Balazy *et al*., 2018; Pinkerton *et al*., 2020; Drago *et al*., 2022). Our results also agree with recent *in situ* imaging data compilation and model output which found that polar waters are dominated by Copepoda whereas in the intertropical waters mixotrophic rhizarians represent a substantial part of the biomass (Biard *et al*., 2016; Drago *et al*., 2022). The median mesozooplankton biomass found in the present study, which varied from 1.3 – 2.9 mgC m^−3^ in the epipelagic (Figure 6 supplementary), is also within the range of biomass reported in the COPEPOD global database (Moriarty and O’Brien, 2013) with values of 0.98 – 3.6 mg C m^−3^.

We calculated zooplankton NBSS slopes derived from UVP5, Multinet samples and the reconstructed NBSS using mode near 1 mm and a linear variation of NBSS with increasing size classes. Despite the significant impact of the previously undersampled rhizarians on NBSS slopes derived from the UVP5 (see next section), especially in tropical and temperate regions, we focus our comparison on global datasets NBSS slopes obtained with nets, as they are the only slopes (for zooplankton-only) reported to our knowledge in previous studies.

The NBSS slopes calculated from the Multinet collection (NBSS_Zmtn) were compared with a global compilation. The comparison showed that the reported values are well within the range of observed values and that they show a contrasting spatial distribution around the globe, consistent with the variability observed between low-productivity and high-productivity systems. The slope of the NBSS is an indicator of the productivity and transfer efficiency of marine ecosystems (Zhou, 2006). In a stable marine ecosystem, the slope should approximately equal to –1 (Sheldon *et al*., 1972). In any given ecosystem, a flatter slope indicates good biomass transfer efficiency from smaller to larger organisms and greater ecosystem stability whereas steeper slopes show the opposite (Sprules and Munawar, 1986). In our study the median slopes of the normalized biovolume size spectra were flatter than −1 (ranging from – 0.57 to – 0.94), which indicates that zooplankton communities in the study area were more characterized by high energy transfer efficiency. The NBSS median slopes from this study were more moderate than those in the North Pacific Ocean, Northwest Pacific Ocean, Northwest Atlantic Ocean, California Current, California Bight and Western Antarctic Peninsula (see Table 3 in (Dai *et al*., 2016)). The median NBSS slopes computed from UVP5 were systematically flatter than the net-collected zooplankton slopes, and also those from the literature.

In our study, steeper slopes were observed in shallow strata than in the mesopelagic. Smaller-sized organisms may prevail in the upper water layer (Ohman and Romagnan, 2016), as they feed on smaller prey such as the phytoplankton and other microzooplankton prey. Being small may be an advantage for escaping predators, but this surface layer is the reproductive layer where all types of larvae prevail. Higher average size, or flatter slopes of organisms in the mesopelagic reflect the relative importance of larger-sized individuals in the deep ocean (Ohman and Romagnan, 2016). Deep ocean environments may favor larger individuals through multiple mechanisms improving their survival. For instance, environmental factors such as lower temperature, less dissolved oxygen, and increased hydrostatic pressure (Childress, 1995) could have decreased the selective pressure for high activity (‘‘predation-mediated selection’’ hypothesis). Also, their diet comprised of diatoms, ciliates, parts of copepods (Harding, 1974; Verity and Paffenhofer, 1996), or fecal pellets through “repackaging” process (Sasaki *et al*., 1988; Lampitt *et al*., 1990) may reduce their metabolic rates and carry high energy contents (Ikeda *et al*., 2006). Larger individuals perform better in deep environments because they can accumulate more energy to survive (Hopcroft *et al*., 2001). However, increased trophic level can also lead to corresponding increases in body size (Romero-Romero *et al*., 2016). The biomass of phytoplankton declines with depth (Yamaguchi *et al*., 2002), so the predator-prey relationship typically shifts from more herbivorous to a more omnivorous or carnivorous ratio (Vinogradov and Tseitlin, 1983). According to (Romero-Romero *et al*., 2016) trophic level is positively correlated with body size. Furthermore, from an individual perspective, deeper living pelagic species use more energy to achieve larger sizes, although they have lower energy concentrations and metabolic rates (Childress *et al*., 1990). As a possible result, a larger body size with greater depth may have been developed as a strategy to increase buoyancy and reduce predator attacks.

### Discrepancy in zooplankton NBSS assessed *in situ* and from net collected samples

Our results demonstrate that assessing zooplankton NBSS and total biomass from samples collected by plankton nets or detected by *in situ* cameras may differ significantly. Many studies have demonstrated that plankton assessment is subjective to the protocol used for collection and analysis (Benfield *et al*., 1998; Harris *et al*., 2000; Wiebe and Benfield, 2003; Remsen *et al*., 2004; Forest *et al*., 2012). However, in the specific case of the Arctic, good agreement was found between net-collected zooplankton and organisms detected *in situ* (Forest *et al*., 2012).

It is well known that nets may discard small organisms as a function of mesh size, and destroy fragile organisms (Remsen *et al*., 2004). *In situ* imaging may be more appropriate for such fragile plankton (Remsen *et al*., 2004; Biard *et al*., 2016) notably in the upper ocean of intertropical regions where large rhizarians (mostly collodarians) predominate. However, a major drawback of *in situ* cameras is the smaller imaged volume, leading to inaccurate analysis of community composition from single casts (Stemmann *et al*., 2008; Picheral *et al*., 2010; Lombard *et al*., 2019). Consistent patterns can however emerge when aggregating multiple UVP5 profiles for each station (in this case with about 10 profiles per station) and/or aggregating results from similar habitats according to latitude and depth as we did. In addition, the taxonomic resolution achieved by *in situ* images is much lower than the taxonomic resolution obtained from net samples scanned with a ZooScan. Due to this methodological biased and to be as accurate as possible, this study focused on organisms larger than >1mm. In this lower size range, many of the particles were identified as marine snow, larvacean houses, diatom rafts, or fecal pellet strings, which we discarded in our analysis as they did not constitute living zooplankton. However, we note that the majority of our unclassified images were of particles smaller than 1 mm ESD, which lacked any resolvable characteristics to support their identification. Thus, it is possible that a large percentage of the small-unidentified particles in the UVP5 dataset were small zooplankton such as early life stages of copepod.

Crustaceans and carnivorous gelatinous were the best groups represented in both UVP and Multinet across all samples. This suggests that UVP5 sampled well crustaceans especially in polar region where their size is bigger, even though the volume imaged was more limited. (Forest *et al*., 2012; Barth and Stone, 2022) observed that the UVP5 did not sample copepods accurately below 1mm ESD. Using copepods as benchmark, since they were abundant and not damaged by the net collection, we showed that for organisms larger than 1mm in ESD, both biomass estimates were generally in agreement in polar regions dominated by crustaceans. Hence in this region, the difference in modal sizes of the zooplankton NBSS was mainly due to difference in the images resolution, rather than by a difference in community composition.

### Novel understanding of pelagic ecology gained by combining *in situ* and net collected data

Our analysis on a global scale mostly in open ocean allow us to draw general conclusions on the relationship between NBSS estimated from *in situ* devices and those estimated from net samples. We showed that in polar regions, where large copepods dominate, both NBSS converge, as also shown by (Forest *et al*., 2012), whereas they showed significant differences in tropical epipelagic layers, as collodarians made up to 70% of the total biovolume (see also (Biard *et al*., 2016)), or in epipelagic layers of temperate ecosystems, as phaeodarians made up to 80% of the total biovolume and 45% in the mesopelagic layers, in agreement with (Biard and Ohman, 2020). In contrast, crustaceans and carnivorous gelatinous organisms always dominated the NBSS derived from net tows.

The average size of the rhizarians found in the UVP5 samples was higher than crustaceans. This suggest that much of plankton biovolume is missed in the net catches taken in inter-tropical regions, as they do not collect efficiently the large rhizarians. However, because of their much lower density, the impact on the zooplankton carbon content may be limited. In upper mesopelagic samples of 0-60°, rhizarians contributed to the total biomass in similar proportion as the contribution of crustaceans, however their contribution decreased with depth. Biomass estimates calculated in this study are consistent with previous estimates, and showed a similar spatial pattern (Biard *et al*., 2016). However, we note that we observed a strong variability of these estimates, depending on the type of the ecosystem and the balance between the fraction of crustaceans (mainly copepods) and rhizarians. It is noticeable that the contribution of large phaeodarians is more important in the mesopelagic layer than in the epipelagic layer. In polar regions, the contribution of rhizarians decrease sharply. The fact that rhizarians are missing in the net lead to underestimation of the energy flow in the food web structure and reduce the efficiency of carbon pump to deep ocean.

The slopes from previous works based on nets samples are steeper than the slopes presented here for tropical and temperate waters and relatively similar in polar waters. The UVP5 slopes are flatter than slopes obtained with other methods. The slopes of the reconstructed NBSS are steeper than the UVP5 and flatter than the Multinet modifying the transfer efficiency estimates from the past studies and the estimation of carbon flow in the food web and in vertical pump. The large variability among slopes of the same latitudinal bands, especially in the epipelagic and upper mesopelagic layers, can be due to the biogeographical conditions and temporal factors (costal/offshore samples, eutrophic/oligotrophic zones and date of cruises) affecting our sampling. In general, the reconstructed slopes were higher than -1, meaning that the transfer efficiency was good or stable. The reconstructed slopes reveal a stable ecosystem in the tropic regions and good transfer of biomass in temperate and polar ecosystems, particulary in the surface layer. Rhizarians well detected by the UVP5 and crustaceans well sampled by the net bring together new pieces to the reconstruction of the new NBSS and slopes.

## Conclusion

We report here, for the first time at global scale, the zooplankton biovolume and carbon biomass estimated in the upper kilometer by the combination of two mature, but incomplete, sampling methods. In theory, *in situ* imaging with UVP5 allows to detect all organisms including fragile ones, albeit in a small sampled volume (several hundreds of liters for each profile), while Multinet combined with ZooScan image analysis samples a large volume (several ten’s of m^−3^ for each sample), but damages fragile organisms. The optimal values measured by both methods are used to reconstruct the ideal zooplankton biovolume and biomass distributions.

Our results showed that the reconstructed NBSS slopes were flatter than the Multinet (- 20%) and steeper than the ones of UVP5 (+7.6%). This suggest that Multinets significantly undersample larges and fragiles organisms because of destruction or avoidance whereas UVP5 is lacking good resolution for the smaller organisms. Large differences between methods were systematically observed in ecosystems dominated by rhizarians, namely the tropical and temperate regions including surface and mesopelagic layers. Thus, the overall gain in polar biomass was relatively small for reconstructed biomass compared to bulk estimates from Multinet (+0.24 mgC/m3 or +4.25%) and high from the UVP5 (+2.0 mgC/m3 or +53%). In contrast, in the tropical and temperate ecosystems, the integrated biomass across size classes between 1 and 8 mm was small in the reconstructed distribution, compared to that of the UVP5 (+0.67 mgC/m3 or +30.44% and +0.74 mgC/m3 or +19.59% respectively) and high from the Multinet (+1.66 mgC/m3 or +136% and +3.4 mgC/m3 or +309% respectively). In the mesopelagic layer there are less difference with reconstructed Biomass when we used UVP5 in comparison to Multinet. These differences suggest that rhizarians, when abundant, have a profound impact on the slope of the NBSS. This biases our ability to use only NBSS calculated from net collected samples as an indicator of the trophic flow of energy, while the high taxonomic resolution of net tows remains important. Lower observed volume and resolution prevents the use of the UVP5 to study mesozooplankton biodiversity. Therefore, with current technologies, both methods should be combined. We recommend the use of only *in situ* imaging technologies when increased resolution and larger field of view are possible. Until then, imaging technologies together with nets samples analysis provides the complete datasets to study ecosystems functioning, using NBSS as a key planktonic variable.

## Acknowledgement

Tara Oceans (which includes both the Tara Oceans and Tara Oceans Polar Circle expeditions) would not exist without the leadership of the Tara Ocean Foundation and the continuous support of 23 institutes (https://oceans.taraexpeditions.org/). The global sampling effort was enabled by countless scientists and crew who sampled aboard the Tara from 2009– 2013, and we thank MERCATOR-CORIOLIS and ACRI-ST for providing daily satellite data during the expeditions. We are also grateful to the countries who graciously granted sampling permission. We thank Agnès b. and Etienne Bourgois, the Prince Albert II de Monaco Foundation, the Veolia Foundation, Region Bretagne, Lorient Agglomeration, Serge Ferrari, Worldcourier, and KAUST for support and commitment. We also thank “Make Our Planet Great Again Team” for the postdoctoral fellow support.

## Data Archiving

Multinet data and UVP5 are published in [PANGAEA](https://www.pangaea.de/submit/). Additional supplementary material was submitted as online Supplementary Material.

## Supplementary

**Table 1 supplementary.**
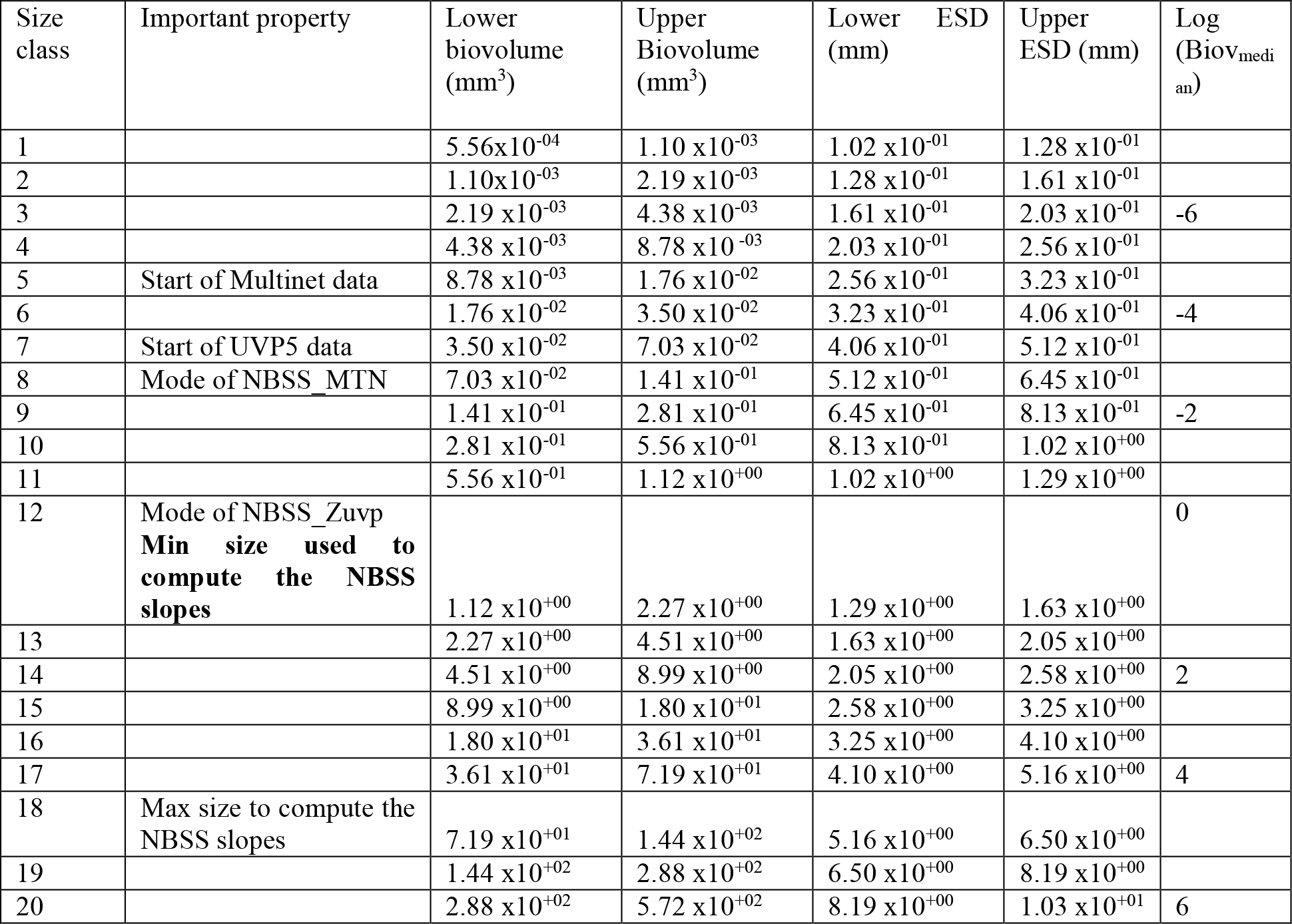
: Lower and upper limits of each size class (biovolume, mm^3^) and corresponding equivalent spherical diameter (ESD, mm) as calculated using the minor and major ellipsoidal axis. The corresponding log(biovolume) reported on Figure 3&4 supplementary is given for reference in the last column

**Table II supplementary.**
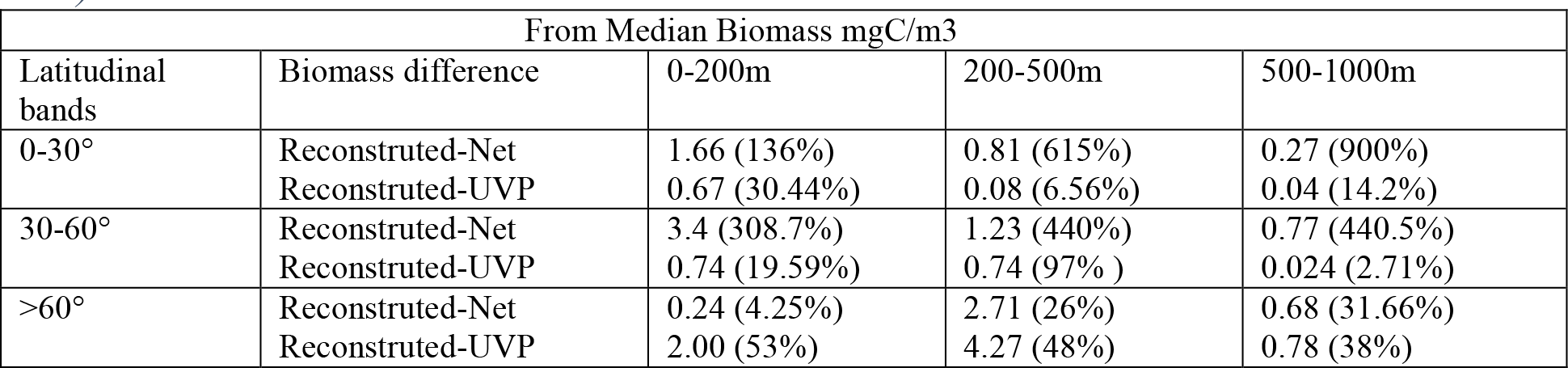
: Biomass difference (reconstructed – Net) and (reconstructed-UVP)

**Figure 1 supplementary:**
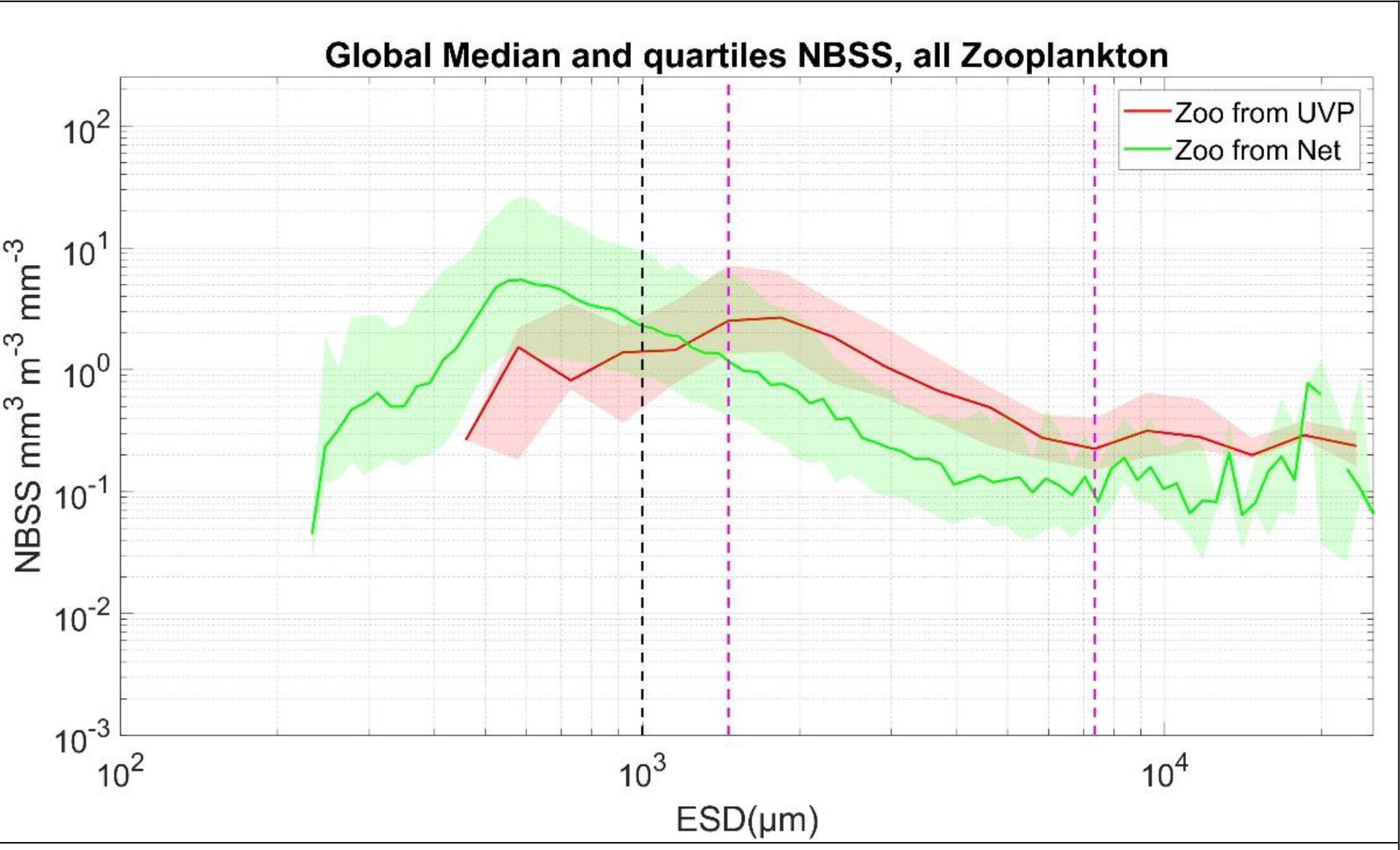
NBSS of all living zooplankton sampled by the UVP5 (red) and nets (green). The shaded area corresponds to the 25 and 75 quartiles around the median of all NBSS observed at the 57 stations shown in Figure 1. The black shaded line corresponds to the 1mm while the marron shaded lines delimit the size range used to extract and to compare the NBSS slopes

**Figure 2 supplementary:**
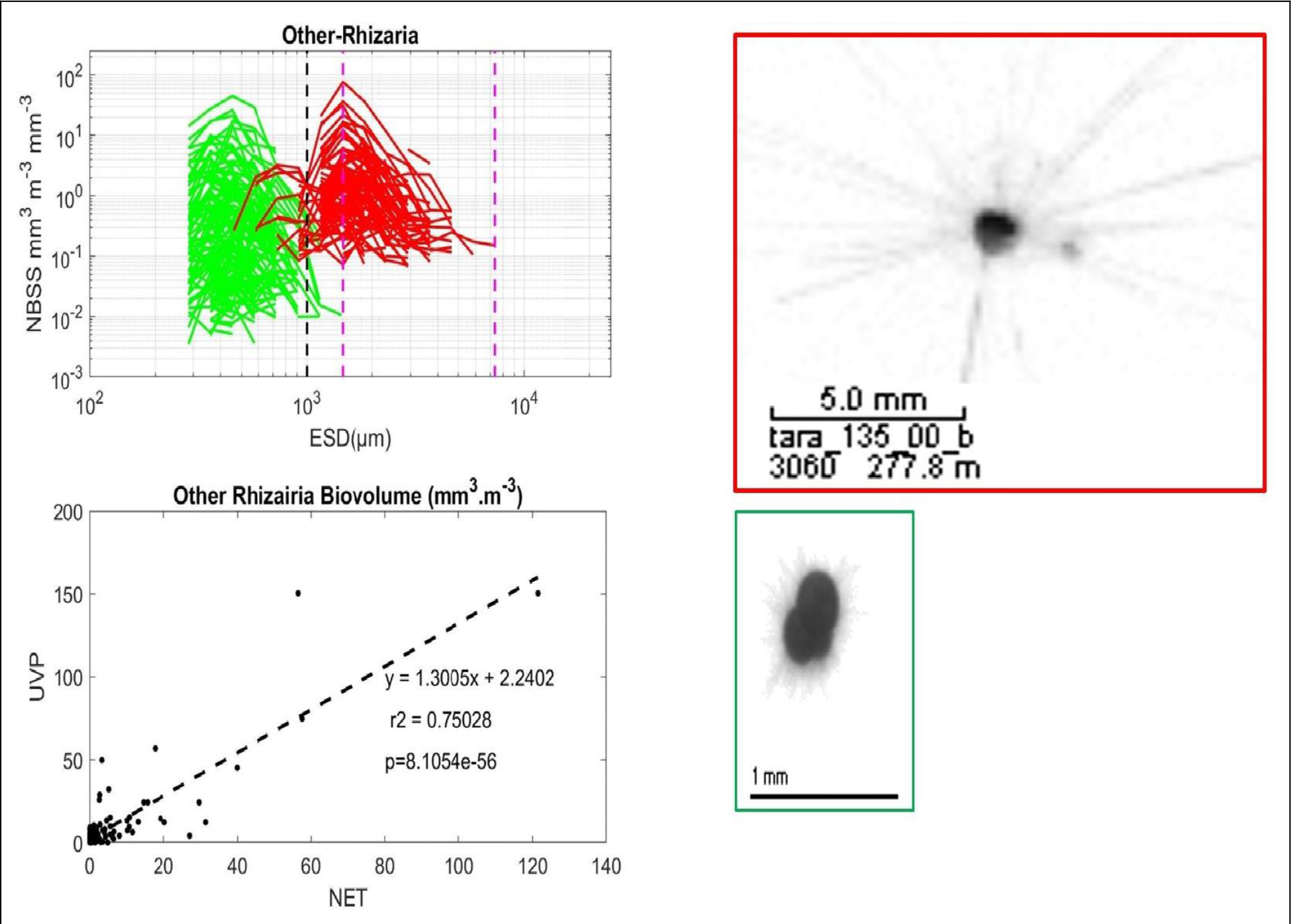
Comparison and correlation of Other Rhizaria from UVP5 (red) and Net (green)

**Figure 3 supplementary :**
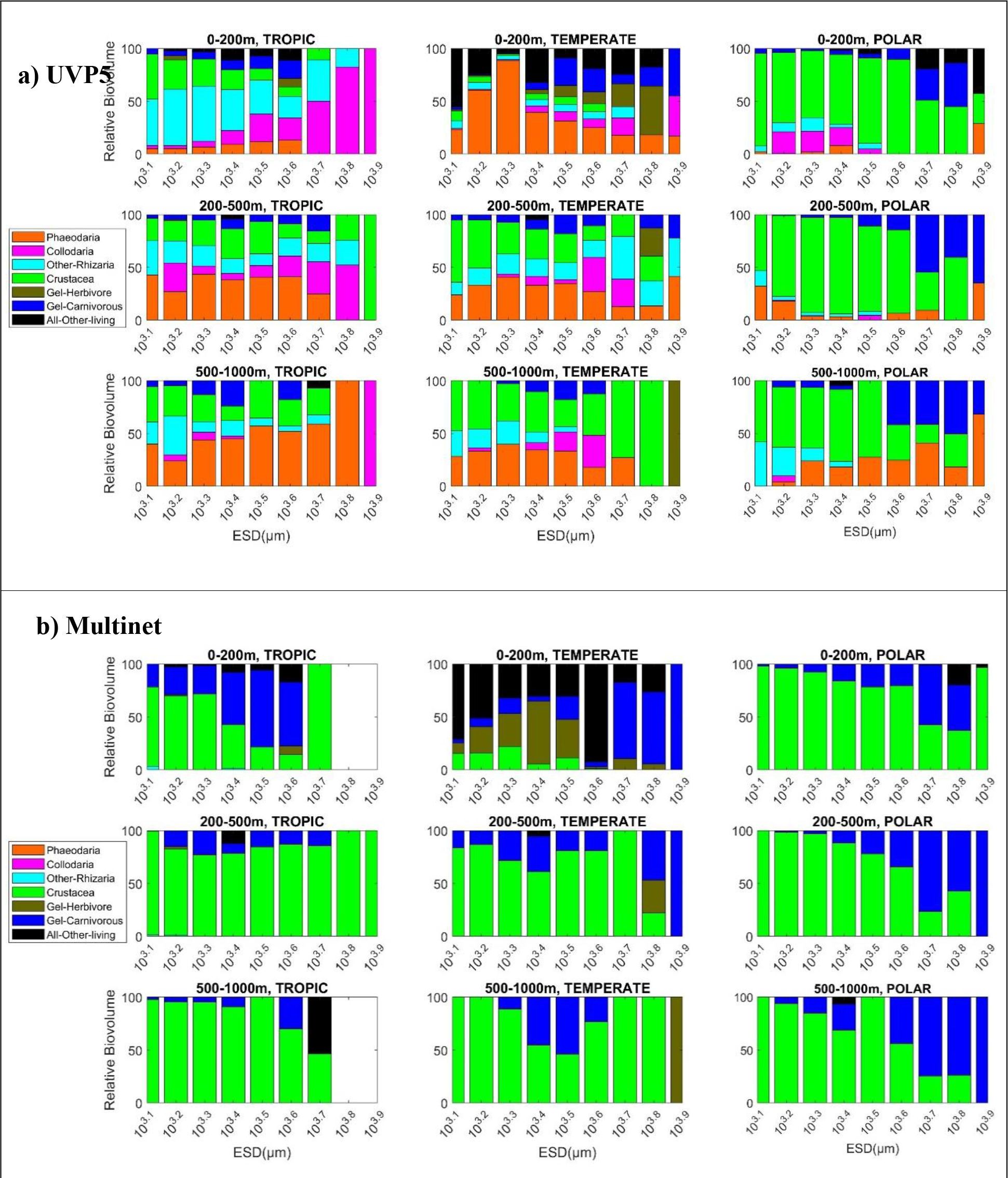
Relative biovolume contributions of different zooplankton taxa detected by the UVP5 (a) or collected with a Multinet (b) in three depth layers (0-200, 200-500 and 500-1000 m) and three latitudinal bands (0°-30°, 30°-60°, >60°) for plankton size from 1 to 8mm.

**Figure 4 supplementary:**
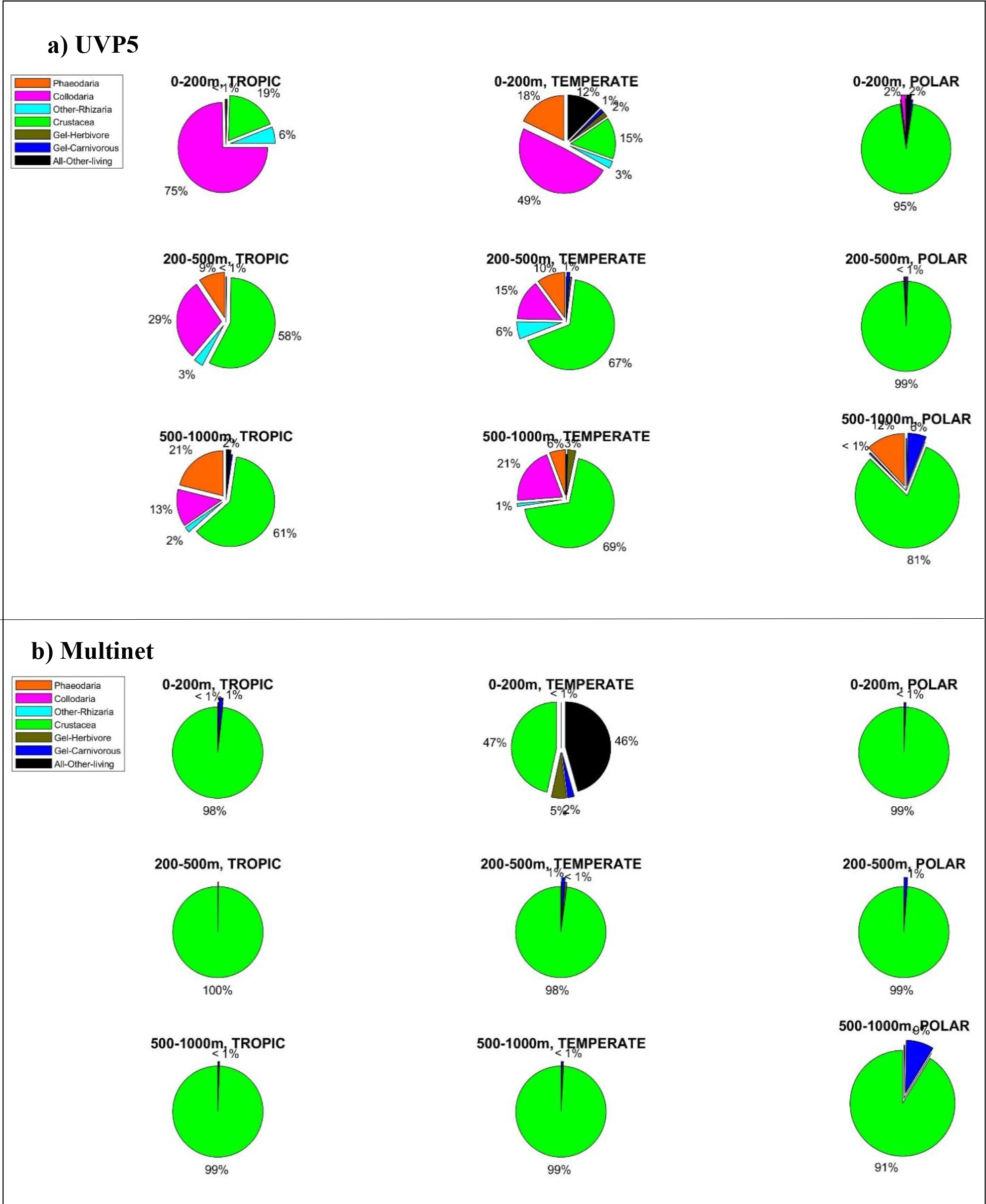
Relative biomass contributions of different zooplankton groups detected by the UVP5 (a) and Multinet (b) in three depth layers (0-200, 200-500 and 500-1000m), and three latitudinal bands (0-30, 30-60°, >60°N/S) for plankton size from 1 to 8mm.

**Figure 5 supplementary:**
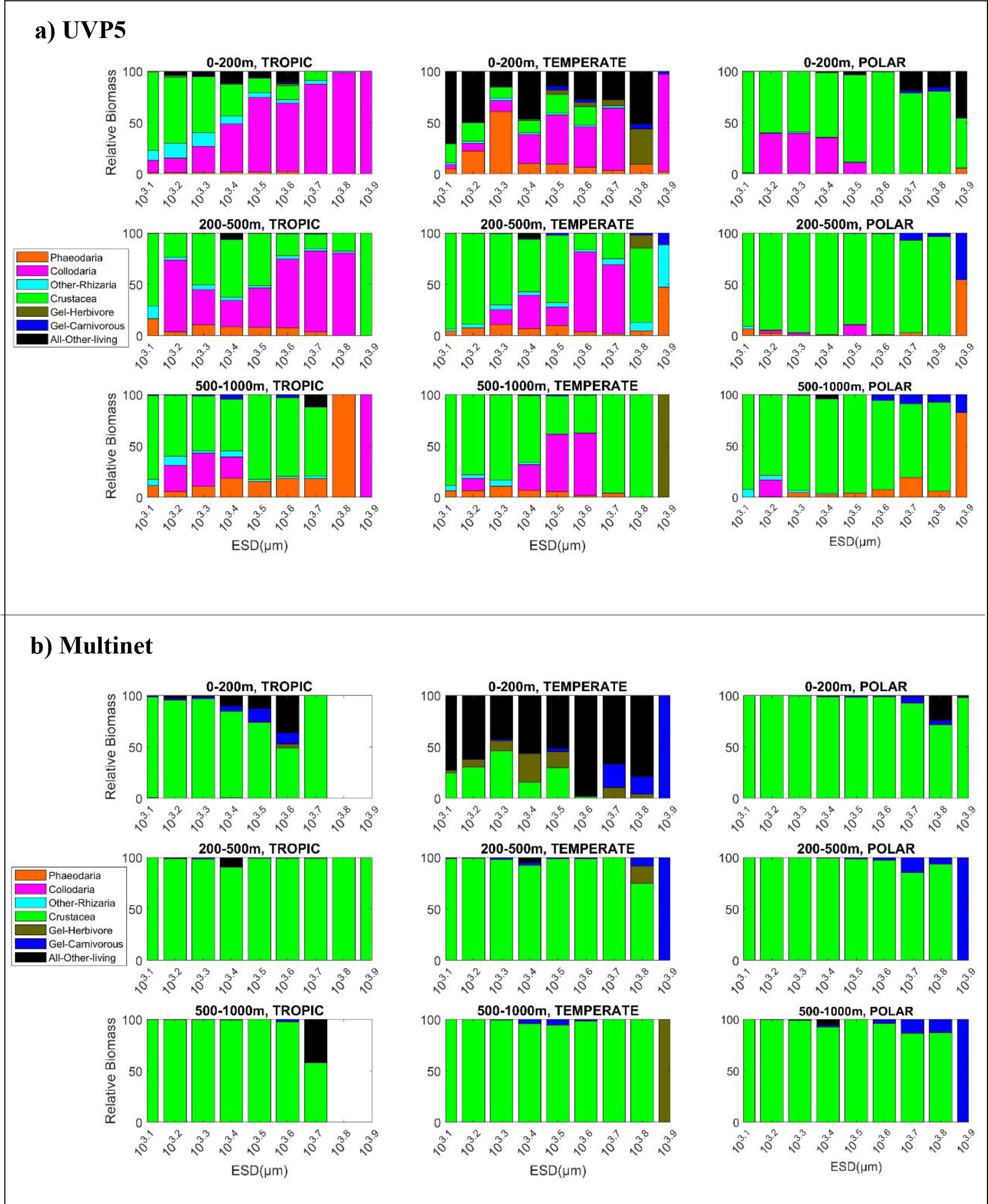
Biomass contributions of different zooplankton taxa sampled with a UVP5 (a) and a Multinet (b) in three depth layers (0-200, 200-500 and 500-1000m), and three latitudinal bands (0-30, 30-60, >60°) for plankton size from 1 to 8mm.

**Figure 6 supplementary.**
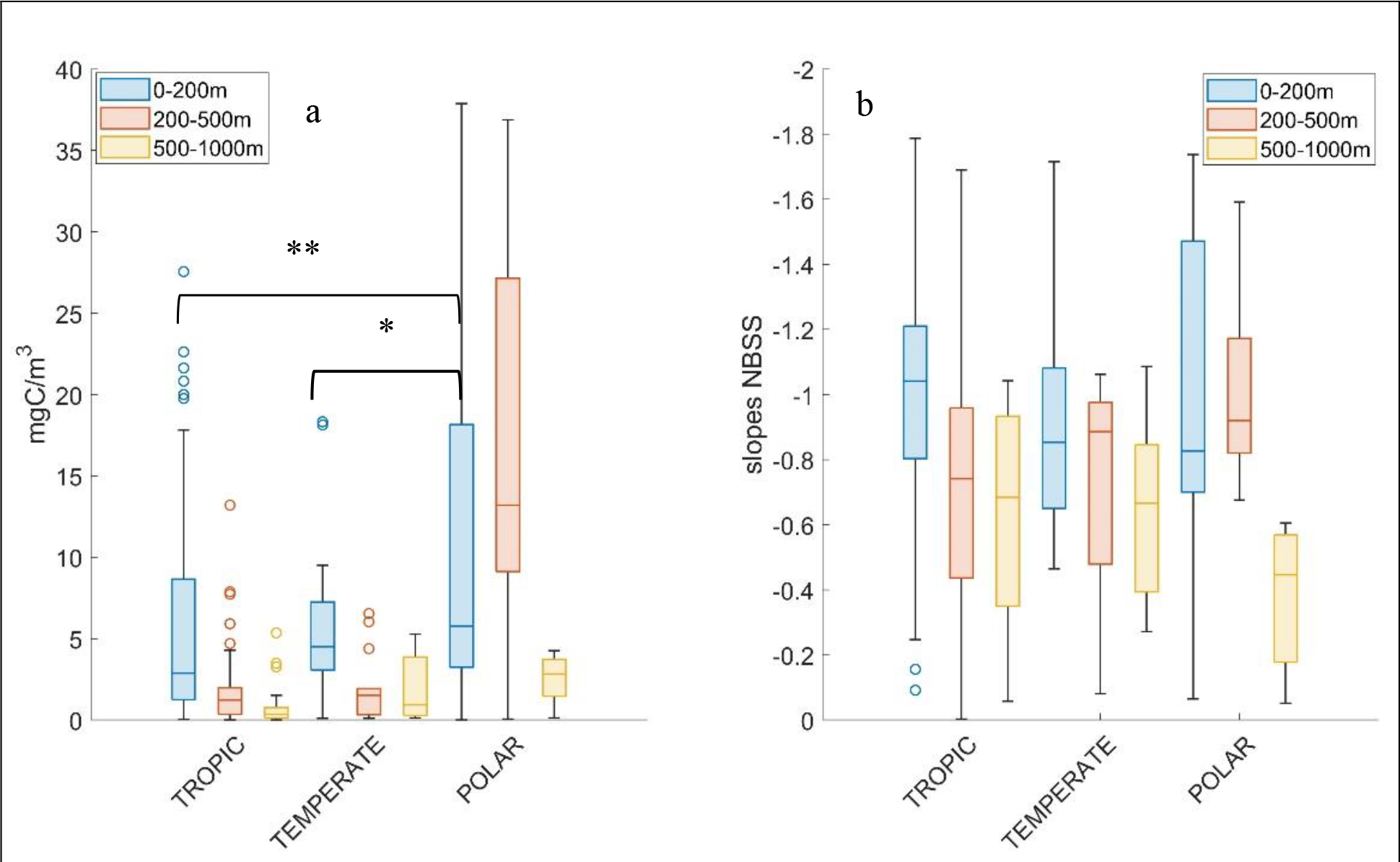
: Distribution of reconstructed biomass (a) and NBSS slopes in Latitudes (b). Statistical Test is ANOVA

**Supplementary Figure 7a.**
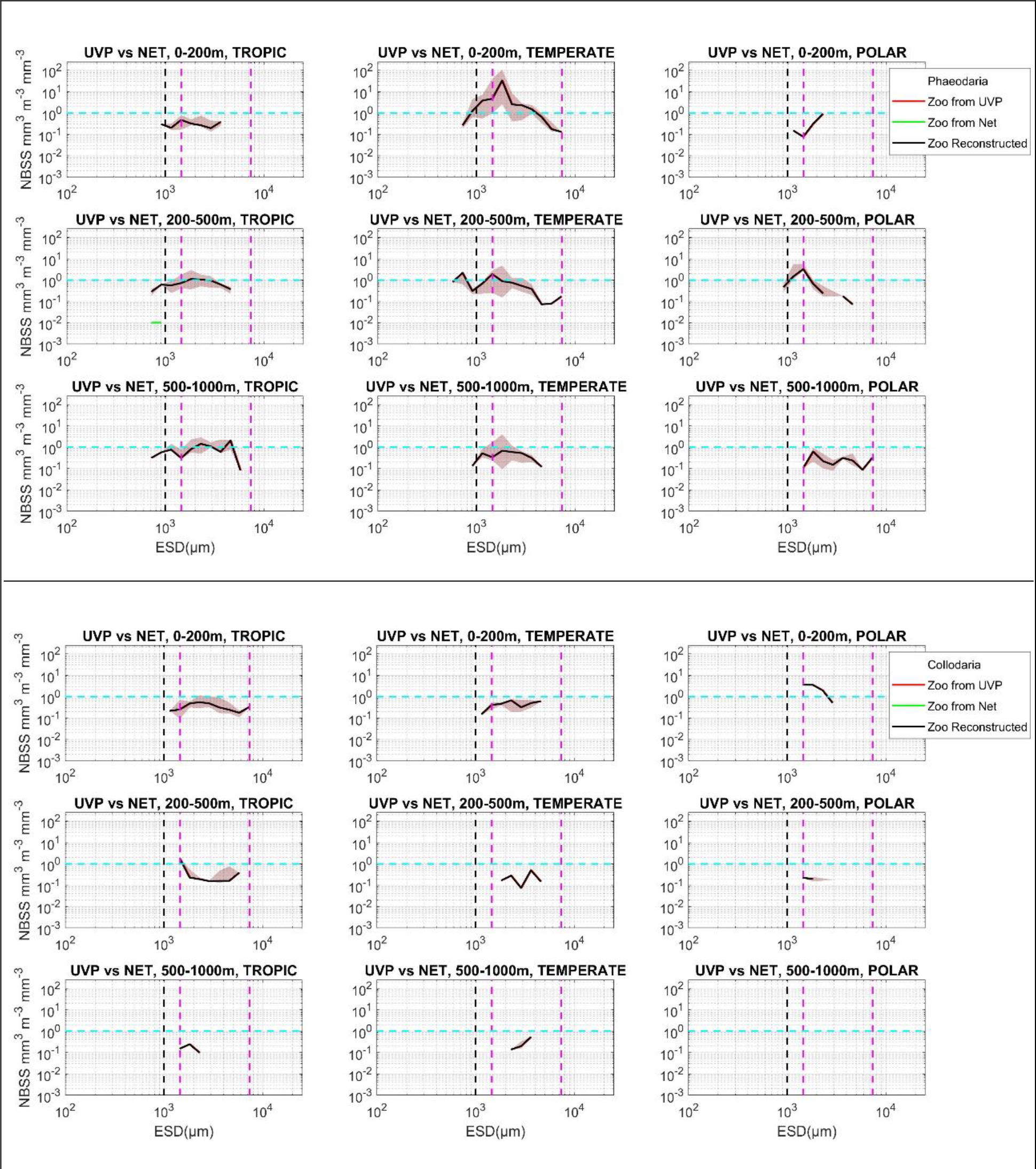
(top:Pheodaria and down:Collodaria) : NBSS maximum selection from Multinet and UVP5

**Supplementary Figure 7b.**
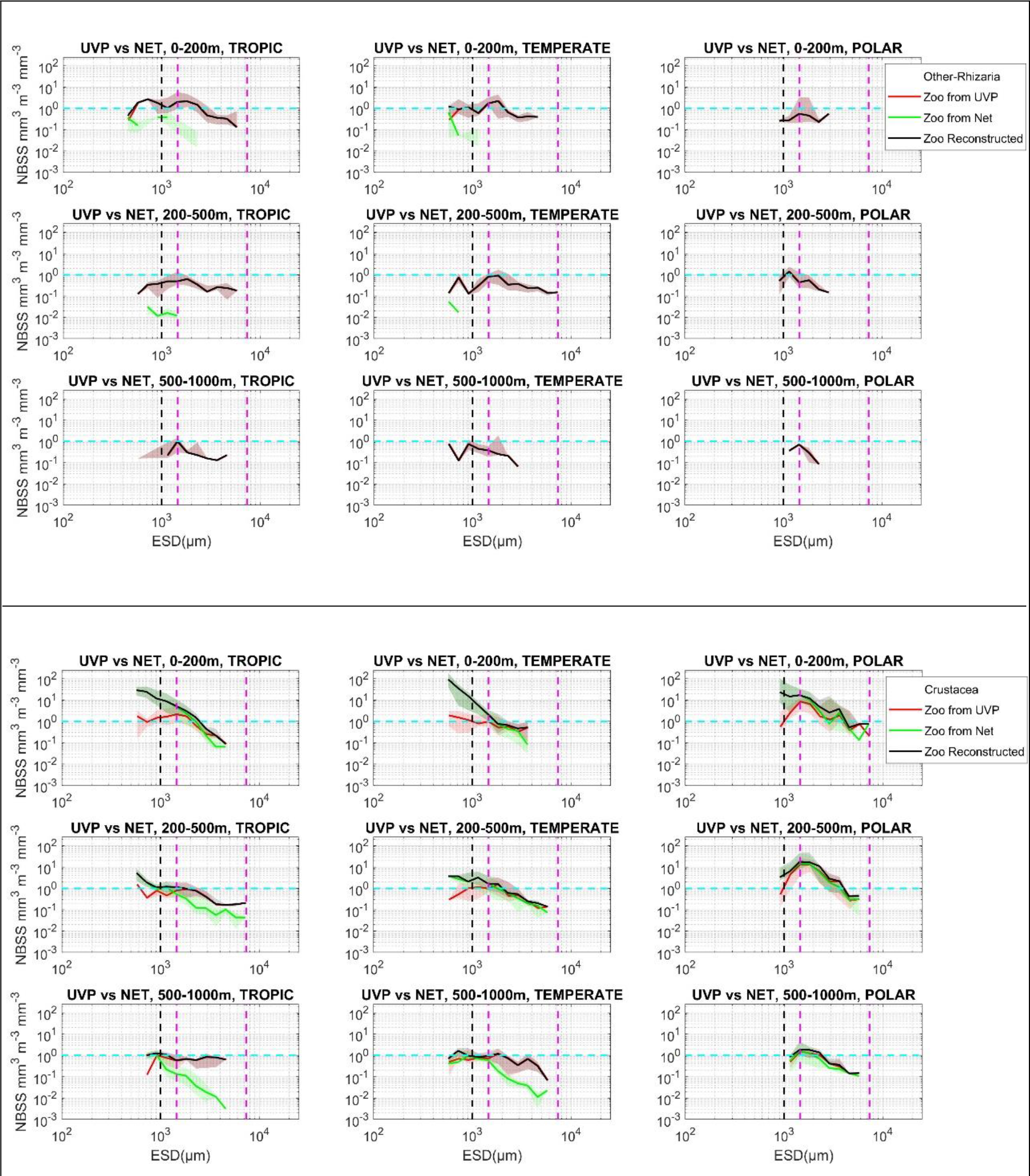
(top:Other Rhizarian and down:Crustacea): NBSS maximum selection from Multinet and UVP5

**Figure 7c supplementary.**
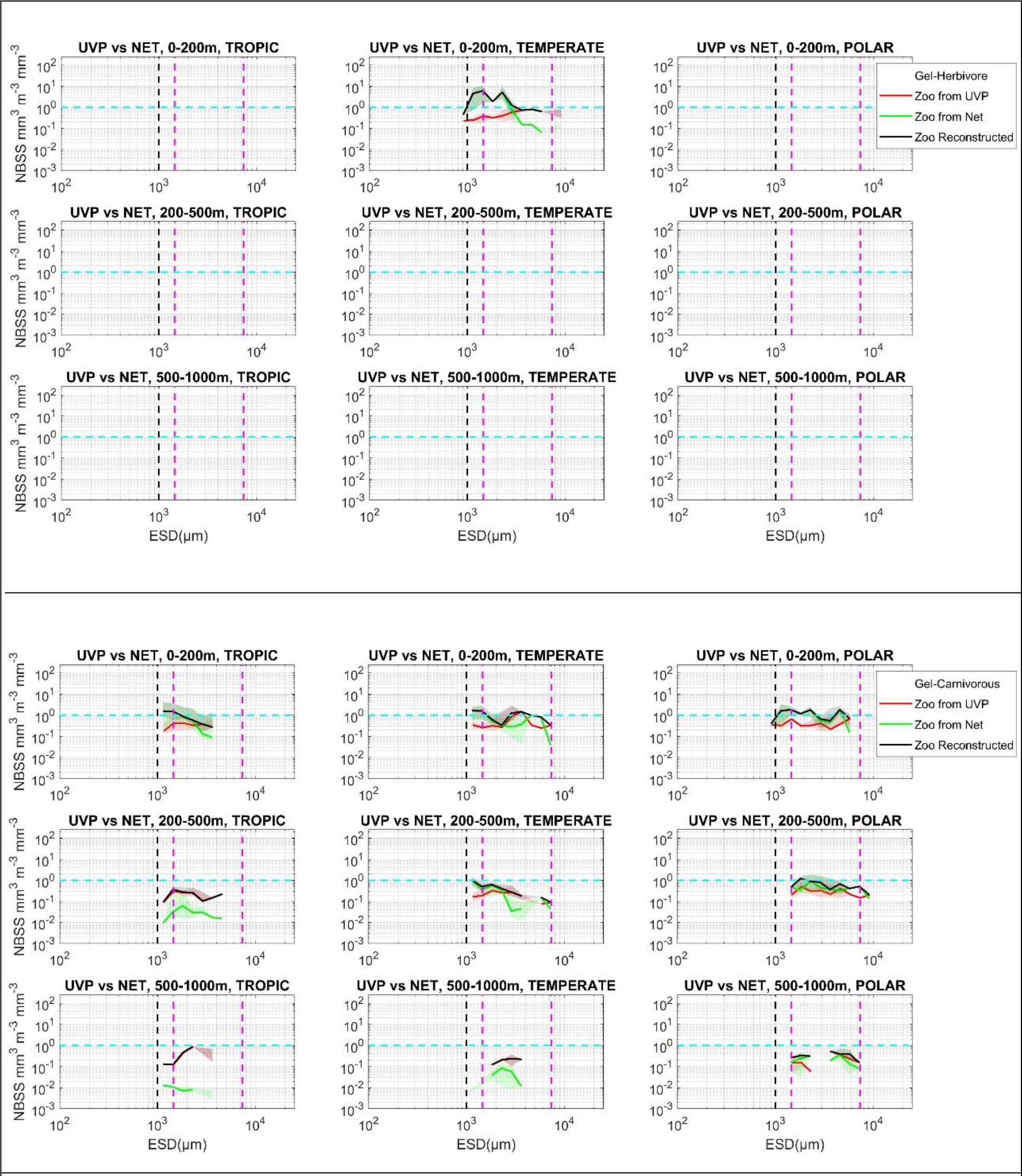
(top:Gelatinous Herbivore and down:Gelatinous Carnivorous): NBSS maximum selection from Multinet and UVP5

**Figure 7d supplementary.**
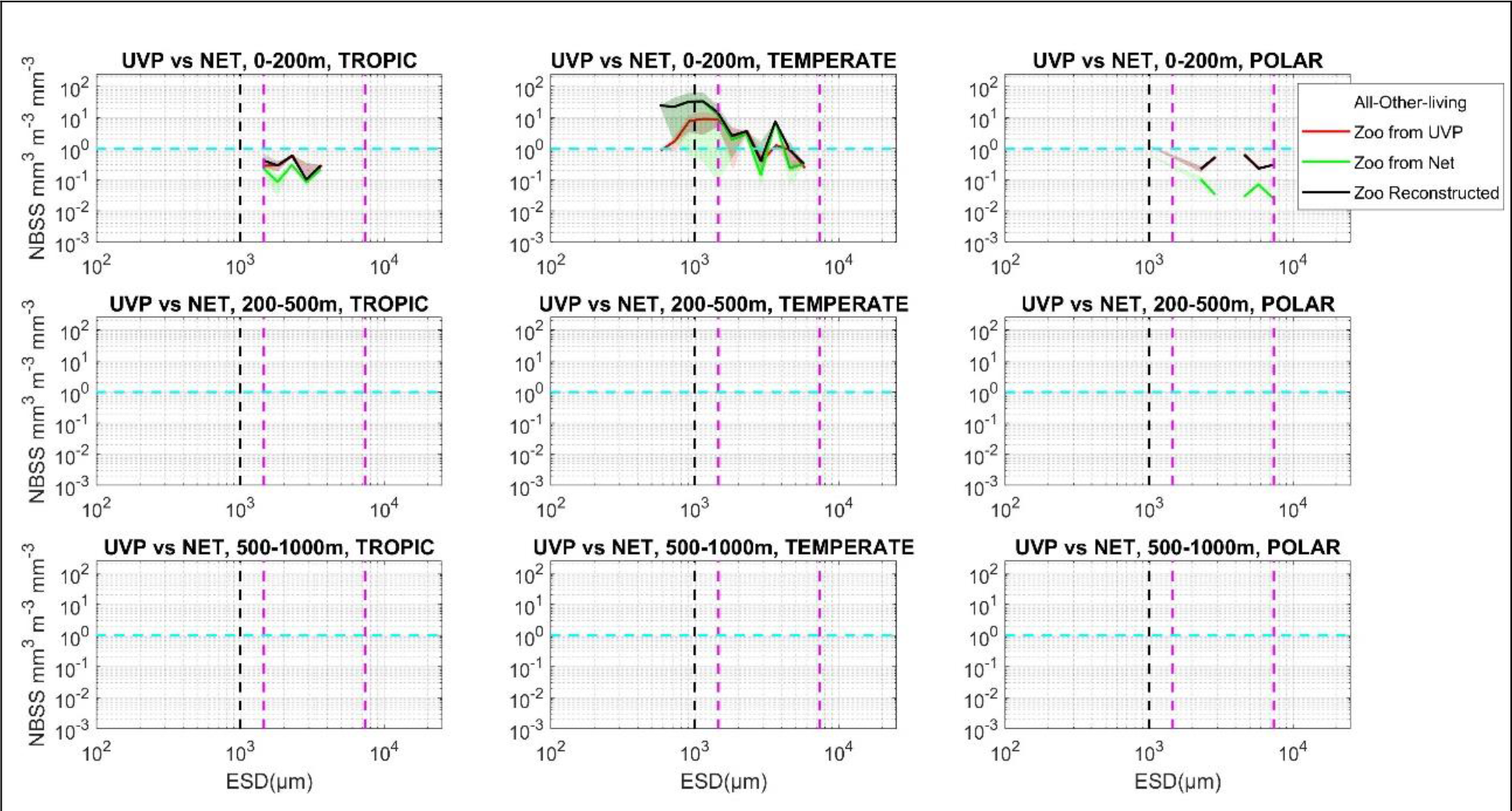
(Other living): NBSS maximum selection from Multinet and UVP5

